# Structural insights into transcription regulation of the global virulence factor PhoP from *Mycobacterium tuberculosis*

**DOI:** 10.1101/2024.05.16.594476

**Authors:** Jing Shi, Qian Song, Zhenzhen Feng, Aijia Wen, Tianyu Liu, Liqiao Xu, Zonghang Ye, Simin Xu, Fei Gao, Liuxiang Xiao, Jiapeng Zhu, Kalyan Das, Guoping Zhao, Yu Feng, Wei Lin

**Affiliations:** School of Medicine, Nanjing University of Chinese Medicine, Nanjing Drum Tower Hospital, Nanjing 210023, China; Department of Biophysics, and Department of Infectious Disease of Sir Run Run Shaw Hospital, Zhejiang University School of Medicine, Hangzhou 310058, China; Rega Institute for Medical Research, Department of Microbiology, Immunology and Transplantation, KU Leuven, Leuven 3000, Belgium; Key Laboratory of Synthetic Biology, CAS Center for Excellence in Molecular Plant Sciences, Chinese Academy of Sciences; Shanghai 200032, China; State Key Laboratory of Bioreactor Engineering, East China University of Science and Technology, Shanghai 200032, China; College of Ecology and Environment, Chengdu University of Technology, Chengdu 610065, China

**Author notes:** Equal contribution (Jing Shi, Qian Song, Zhenzhen Feng, Aijia Wen).

**Keywords:** *Mycobacterium tuberculosis*, PhoP, cryo-EM structure, PhoP-TACs, PhoP-TRC

## Abstract

*Mycobacterium tuberculosis* (*Mtb*), remaining as the leading cause of the worldwide threat Tuberculosis, relies heavily on its transcriptional reprogramming of diverse stress genes to swiftly adapt to adverse environments and ensure infections. The global virulence factor PhoP plays a pivotal role in coordinating transcription activation or repression of the essential phosphate-nitrogen metabolic remodeling genes. However, what defines PhoP to deferentially act as an activator or a repressor remains largely unexplored. Here, we determine one cryo-EM structure of *Mtb* RNAP-promoter open complex, three cryo-EM structures of PhoP-dependent transcription activation complexes (PhoP-TACs) consisting of *Mtb* RNA polymerase (RNAP), different number of PhoP molecules binding to different types of well-characterized consensus promoters, and one cryo-EM structure of *Mtb* PhoP-dependent transcription repression complex (PhoP-TRC) comprising of *Mtb* RNAP, PhoP, the nitrogen metabolism regulator GlnR and their co-regulated promoter. Structural comparisons reveal phosphorylation of PhoP is required for stabilization of PhoP-TACs, PhoP specifically recognizes promoters as novel tandem dimers and recruits RNAP through extensively interacting with its conserved β flap and σ^A^R4 domains. Strikingly, the distinct promoter spacer length and PhoP-GlnR interactions in PhoP-TRC constrain the upstream DNA into a distinct topology and retain PhoP in a novel ‘dragging repression mode’. Collectively, these data highlight the dual regulatory mechanisms of PhoP-dependent transcription regulation in governing stress adaptation. These findings provide structural basis for developing potential anti-tuberculosis drugs and/or interventions.

## Introduction

*Mycobacterium tuberculosis* (*Mtb*), as the leading causative pathogen of the worldwide disease Tuberculosis (TB), still infects almost one third of the human population and kills two million people every year^1^. Emergence of drug-resistant and multidrug-resistant *Mtb* strains makes treating TB increasingly challenging. Nowadays, it is particularly urgent to explore the molecular mechanisms underlying infections, and to develop novel drugs and target interventions^2^. Being an intracellular pathogen, *Mtb* has evolved highly complex transcriptional reprogramming network to quickly adapt to the constantly changing micro-environmental stresses (nutrient deprivation, hypoxia, low pH, *et al*) within host cells^3,4,5^. The most commonly and efficiently adopted strategy is precisely coordinating expression of diverse essential metabolism genes by the global transcription regulators^6,7^.

Phosphorus is one of the essential nutrients required for synthesis of biological macromolecules. Thereby stringent regulation of phosphate metabolism is crucial for all life beings. During infection, *Mtb* usually endures severe stresses of phosphate deficiency by host immune cells. This crisis is supposed to be salvaged by finely activating transcription initiation of assimilation genes involved in inorganic and organic phosphate metabolism (the high-affinity phosphate-specific transport systems genes *pstSCAB*, inorganic phosphate transport genes *pit*, the global phosphate starvation stress response modulator PhoU encoding gene *phoU,* and so on)^8,9^, or repressing transcription of relevant dissimilation genes or the other metabolic genes, such as repressing the nitrogen metabolism gene *amtB* via cross-talking with the global nitrogen metabolism regulator GlnR^10,11,12^. Transcription regulation of these intricate phosphate-nitrogen metabolic remodeling is elegantly coordinated by a global virulence factor PhoP, which is highly conserved in *actinobacteria* and plays important regulatory roles in maintaining the aforementioned cellular phosphate homeostasis by serving as one essential component of the widespread two-component systems (TCSs) involved in signal transduction^6^. Even more intriguing, PhoP is also evidenced to be involved in regulation of virulence, low pH, hypoxia, respiration, cell differentiation, and antibiotic biosynthesis^4,5^. As supports the pivotal role of PhoP as a key virulence factor in remodeling the transcriptional network of *Mtb* when facing adverse cellular stimuli, further exploration on its molecular determinants may have high implications in developing new target drugs or interventions.

As a paradigm of the enormous OmpR/PhoB subfamily transcription regulators, PhoP also encompasses a conserved N-terminal receiver domain (REC) which can be phosphorylated by the sensor kinase PhoR *in vivo* or acetyl phosphate (AcP) *in vitro*, and thereby enhances the DNA binding affinity of the C-terminal DNA-binding domain (DBD), nevertheless, the DNA binding activity is observed to be phosphorylation-independent^13,14,15,16^. Generally, the specific sequences recognized by PhoP (named PHO boxes) usually contain direct repeat units (DRus), each of which is constituted by two conserved PhoP binding sites (a site and b site) separated by several bases (**Fig. 1**)^15,17,18^. This implies that PhoP may function as possible dimer or dimers to recognize the repeated PHO boxes like the other OmpR/PhoB subfamily transcription regulators^7,13,19,20,21,22^. As is known, bacterial transcription initiation is predominantly driven by the highly conserved multi-subunit RNA polymerase (α2ββ’ωσ) harboring the promoter specific factor σ, and elaborately regulated by various transcription regulators when encountering different environmental stresses^23,24,25,26,27^. However, what defines PhoP to deferentially act as a transcription activator or serve as a transcription repressor remains poorly understood.

**Fig. 1.**
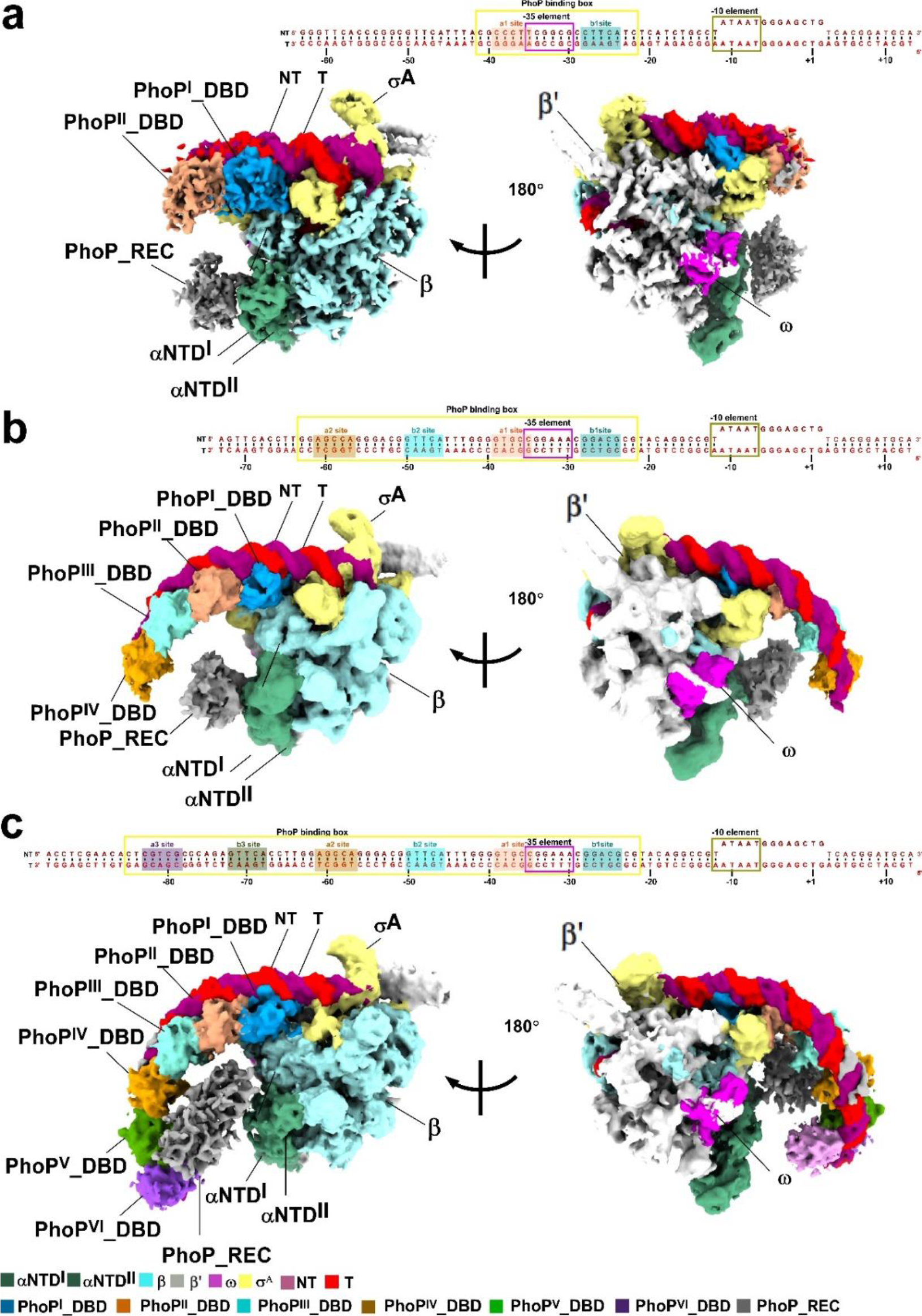
The overall structures of *Mtb* PhoP-TAC. **a,** DNA scaffold used in structure determination of *Mtb* 2PhoP-TAC (top panel) and two views of the cryo-EM density map of *Mtb* 2PhoP-TAC (bottom panel). **b,** DNA scaffold used in structure determination of *Mtb* 4PhoP-TAC (top panel) and two views of the cryo-EM density map of *Mtb* 4PhoP-TAC (bottom panel). **c,** DNA scaffold used in structure determination of *Mtb* 6PhoP-TAC (top panel) and two views of the cryo-EM density map of *Mtb* 6PhoP-TAC (bottom panel). According to the structures, the PhoP binding box, -35 element, -10 element is boxed with yellow, violet, and brown colors, respectively. a1 site, b1 site, a2 site, b2 site, a3 site and b3 site of the PhoP binding box are shaded with light orange, dark cyan, brown, cyan, purple, and dark green, respectively. The EM density maps are colored as indicated in the color key. NT, non-template-strand promoter DNA; T, template-strand promoter DNA.

Here, we determine three cryo-EM structures of PhoP-dependent transcription activation complexes (2PhoP-TAC, 4PhoP-TAC, 6PhoP-TAC) consisting of *Mtb* RNA polymerase (RNAP), *Mtb* PhoP protein, and the well-characterized consensus promoters including different number of PHO boxes, respectively. Structural comparisons with the simultaneously obtained *Mtb* RNAP-promoter open complex (RPo) reveals that phosphorylation of PhoP is required for formation and stabilization of the PhoP-TACs. In PhoP-TACs, PhoP specifically recognizes promoter PHO boxes as dimer or novel tandem dimers (tetramer/hexamer), and makes abundant protein-protein interactions with the conserved domains (β flap and σ^A^R4) of RNAP to stabilize the complexes. Moreover, we also resolve one cryo-EM structure of PhoP-dependent transcription repression complex (PhoP-TRC) which contains *Mtb* RNAP, PhoP, the nitrogen metabolism regulator GlnR, and the nitrogen assimilation promoter *amtB*. Notably, PhoP is constrained in a novel ‘dragging repression mode’ in PhoP-TRC, which is strengthened by the longer upstream spacer length. Though the structure defines similar protein-DNA interactions to those occurred in PhoP-TACs and the recently reported GlnR-dependent transcription complex (GlnR-TAC), it reveals different protein-protein interactions between PhoP and GlnR. These structural and supporting biochemical data collectively reveal novel transcription regulation mechanisms of the global regulator PhoP, which finely keep balance on the physiological phosphate-nitrogen homeostasis under nutrition restriction conditions. These comprehensively enrich our understanding on the crucial regulatory roles of the PhoP-like OmpR/PhoB subfamily transcription regulators on governing transcription plasticity and restoring pathogenic adaptation. This study uncovers new structural targets for exploring promising anti-TB drugs and/or interventions.

## Results

### Overall structure of PhoP-TAC and PhoP-TRC

To dissect the structural basis of PhoP-dependent transcription regulation, we reconstituted the PhoP-dependent transcription activation complexes (PhoP-TACs) by using the purified *Mtb* RNAP, *Mtb* PhoP protein, and the characterized phosphate metabolism promoters with different number of the PHO boxes (**Fig. 1** and **Supplementary Figs. 1, 2**)^8,18,28^. The high-affinity phosphate specific transporter promoter *pst* contains one PHO box with the a1 site and b1 site that overlap with the promoter -35 element (**Fig. 1a**). The global phosphate starvation response modulator promoter *phoU4* includes two PHO boxes with the upstream a2-b2 sites and the downstream a1-b1 sites (**Fig. 1b**). When extends the *phoU4* scaffold by 27 base pairs (bp), it forms a longer *phoU6* scaffold which includes three PHO boxes with the upstream a3-b3 sites, the middle a2-b2 sites, and the downstream a1-b1 sites (**Fig. 1c** and **Supplementary Figs. 1**). By analogy, the PhoP-dependent transcription repression complex (PhoP-TRC) was assembled by incubating the *Mtb* RNAP, PhoP protein, GlnR protein, the PhoP-GlnR co-regulated nitrogen assimilation promoter *amtB* which was previously confirmed to be repressed by combination of PhoP and GlnR^10,12^, and then purified to high homogeneity as well (**Supplementary Figs. 1** and 2). Meanwhile, the downstream part of each promoter is annealed to carry a consensus –10 element, a 13-nucleotide transcription bubble, and an 11-bp downstream double stranded DNA as reported^7,29,30^. To improve the DNA binding affinity, the PhoP proteins used to assemble the PhoP-TACs or PhoP-TRC were all phosphorylated with AcP before incubation with each DNA scaffold and the RNAP holoenzyme^15,16^.

As expected, SDS-PAGE analysis of each complex purified by size-exclusion chromatography showed appearance of each assembled component, in despite of different ratio of PhoP proteins and GlnR proteins (**Supplementary Figs. 2**a-2e). Consistently, the *in vitro* transcription assays with PhoP target promoter DNA fused with a fragment of *mango* sequence showed higher transcription activity of PhoP on the promoter *phoU6* (containing 6 PhoP binding sites) than on the promoter *phoU2* (containing 2 PhoP binding sites) and the promoter *phoU4* (containing 4 PhoP binding sites) (**Supplementary Figs. 2**f), indicative of a higher transcription efficiency for 6PhoP involved. Meanwhile, addition of PhoP impaired about half of the GlnR-dependent transcription activities on the PhoP-GlnR co-regulated promoter *amtB* (**Supplementary Figs. 2**g). These observations indicate that all of the proteins purified are enzymatically active, and PhoP plays dual transcription regulatory roles in the phosphate- nitrogen metabolism genes as previously identified^4,9,12^. The PhoP-dependent transcription complexes obtained showed promising profiles for further cryo-EM analysis.

After repeated attempts in optimizing the samples and data collection, we finally obtained one dataset of RPo just containing *Mtb* RNAP and promoter DNA at a nominal resolution of 3.40 Å, three datasets of PhoP-TAC with different number of PhoP molecules included at nominal resolutions of 3.20 Å, 4.50 Å, and 3.20 Å, and one dataset of PhoP-TRC encompassing both PhoP and GlnR proteins at a nominal resolution of 3.68 Å (**Figs. 1-4**, **Table 1**, and **Supplementary Figs. 3**-12). In each of the cryo-EM map, it shows clear densities for the major and minor grooves of the downstream promoter DNA as well as each subunit of RNAP, which are sufficient to well fit the model of *Mtb* RPo (PDB ID: 6VVY) into them (**Supplementary Figs. 13**-19)^31^. Compared with the main body of RNAP that exhibits a local resolution at ∼ 3.0–4.5 Å, the peripheral maps of αCTD and PhoP molecules interacting with the upstream promoter DNA are likely more flexible (**Supplementary Figs. 8**c, 9c, 10c, and **11c**). Surprisingly, our recently resolved model of *Mtb* GlnR-TAC (PDB ID: 8HIH) can be fitted into most densities of PhoP-TRC^7^. In contrast, the unambiguous densities for PhoP engagement at the upstream DNA and the resulting significant DNA kink suggest PhoP is stabilized in a novel transcription repression mode in PhoP-TRC (**Fig. 4** and **Supplementary Figs. 7**, 12 and 19).

**Table 1.**
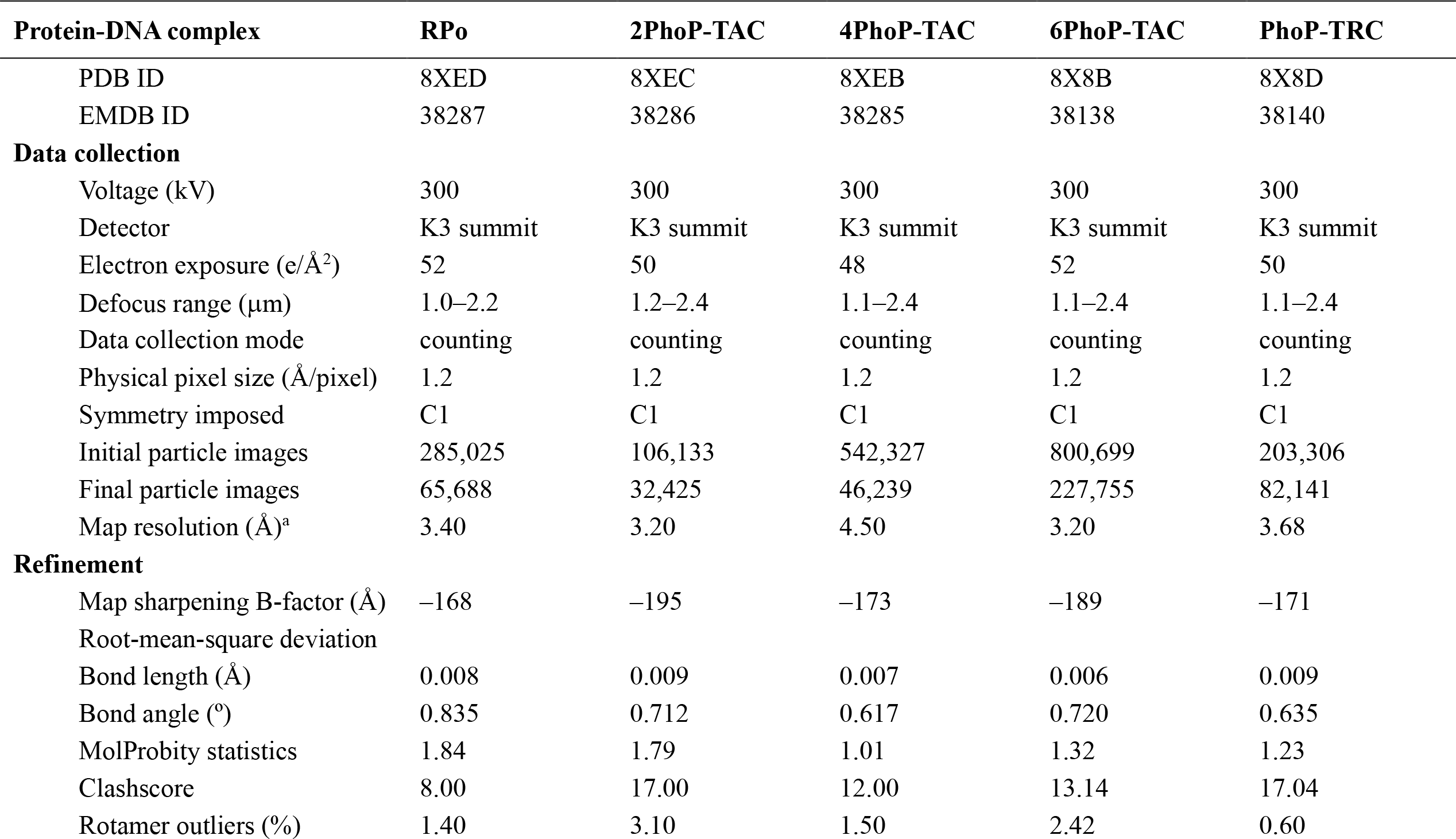
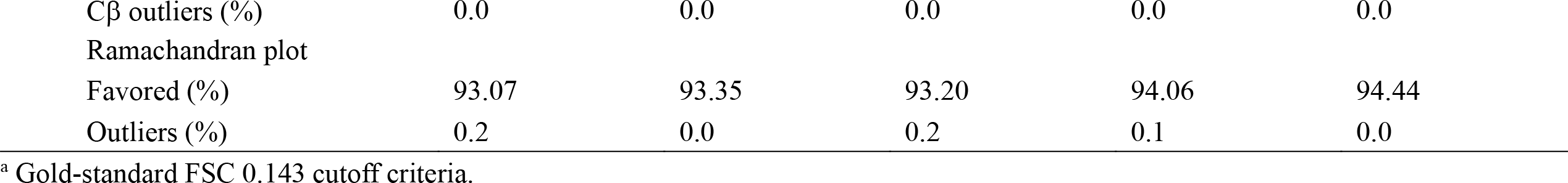
Single particle cryo-EM data collection, processing, and model building for *M. tuberculosis* RPo, 2PhoP-TAC, 4PhoP-TAC, 6PhoP-TAC, and PhoP-TRC.

| Protein-DNA complex | RPo | 2PhoP-TAC | 4PhoP-TAC | 6PhoP-TAC | PhoP-TRC |
| --- | --- | --- | --- | --- | --- |
| PDB ID | 8XED | 8XEC | 8XEB | 8X8B | 8X8D |
| EMDB ID | 38287 | 38286 | 38285 | 38138 | 38140 |
| **Data collection** |  |  |  |  |  |
| Voltage (kV) | 300 | 300 | 300 | 300 | 300 |
| Detector | K3 summit | K3 summit | K3 summit | K3 summit | K3 summit |
| Electron exposure (e/Å<sup>2</sup>) | 52 | 50 | 48 | 52 | 50 |
| Defocus range (μm) | 1.0–2.2 | 1.2–2.4 | 1.1–2.4 | 1.1–2.4 | 1.1–2.4 |
| Data collection mode | counting | counting | counting | counting | counting |
| Physical pixel size (Å/pixel) | 1.2 | 1.2 | 1.2 | 1.2 | 1.2 |
| Symmetry imposed | C1 | C1 | C1 | C1 | C1 |
| Initial particle images | 285,025 | 106,133 | 542,327 | 800,699 | 203,306 |
| Final particle images | 65,688 | 32,425 | 46,239 | 227,755 | 82,141 |
| Map resolution (Å)<sup>a</sup> | 3.40 | 3.20 | 4.50 | 3.20 | 3.68 |
| **Refinement** |  |  |  |  |  |
| Map sharpening B-factor (Å) | −168 | −195 | −173 | −189 | −171 |
| Root-mean-square deviation |  |  |  |  |  |
| Bond length (Å) | 0.008 | 0.009 | 0.007 | 0.006 | 0.009 |
| Bond angle (°) | 0.835 | 0.712 | 0.617 | 0.720 | 0.635 |
| MolProbity statistics | 1.84 | 1.79 | 1.01 | 1.32 | 1.23 |
| Clashscore | 8.00 | 17.00 | 12.00 | 13.14 | 17.04 |
| Rotamer outliers (%) | 1.40 | 3.10 | 1.50 | 2.42 | 0.60 |
| Cβ outliers (%) | 0.0 | 0.0 | 0.0 | 0.0 | 0.0 |
| Ramachandran plot |  |  |  |  |  |
| Favored (%) | 93.07 | 93.35 | 93.20 | 94.06 | 94.44 |
| Outliers (%) | 0.2 | 0.0 | 0.2 | 0.1 | 0.0 |

<sup>a</sup> Gold-standard FSC 0.143 cutoff criteria.

Though the densities for PhoP-REC and GlnR-REC are incomplete in the structures, the ones next to RNAP αNTD give us new hints for their potential regulatory roles in accumulating stabilization of PhoP-TRC.

To get a better understanding, we take the RPo that includes no PhoP protein as a negative control, and designate the high resolution PhoP-TACs containing two PhoP molecules, four PhoP molecules, six PhoP molecules as 2PhoP-TAC, 4PhoP-TAC, and 6PhoP-TAC, respectively. Since the protein-DNA, protein-protein interactions displayed in 2PhoP-TAC and 4PhoP-TAC are highly similar to those involved in 6PhoP-TAC, we choose 6PhoP-TAC to describe the structural basis of PhoP-dependent transcription activation, and use PhoP-TRC to uncover the structural basis of PhoP-dependent transcription repression.

### Phosphorylation of PhoP facilitates stabilization of PhoP-dependent transcription regulation complexes

As evidenced, phosphorylation of response regulators plays an important role in bacterial adaptation through activating or repressing gene expression, especially for the well characterized TCSs which widely mediate cellular signal transduction. Like the other reported OmpR*/*PhoB subfamily transcription factors, phosphorylation of PhoP was demonstrated to induce dimerization of the REC domains, and thereby strengthens the specific promoter binding affinity of the DBDs^19,20,21^. In line with this, we only obtained a structure of *Mtb* RPo when we assembled the PhoP-dependent transcription complexes with the wild-type PhoP proteins not being phosphorylated. While it tends to be much easier to classify and resolve the functional structures of PhoP-TACs or PhoP-TRC when using the AcP-phosphorylated PhoP proteins instead (**Figs. 1-4**, and **Supplementary Figs. 2**-13). These indicate phosphorylation of PhoP is also required for formation and stabilization of the PhoP-dependent transcription regulation complexes. Therefore, each of the other PhoP-dependent transcription complexes was also prepared by the AcP-phosphorylated PhoP proteins which were referred to as PhoP in the following text. It is most probably triggered by the synergistic conformation changes of the REC and DBD domains as reported, which further evoke dimerization and enhance cooperative binding of the tandem PhoP molecules to the PHO boxes as observed in the cryo-EM structures of PhoP-TACs and PhoP-TRC.

### Cryo-EM analysis of different PhoP-TACs reveals the structural basis of PhoP functioning as a global transcription activator

By comparison with *Mtb* RPo and different PhoP-TACs, the hallmarks of PhoP-dependent transcription activation are clearly outlined: PhoP specifically recognize promoter PHO boxes as distinct tandem dimer/tetramer/hexamer, and make abundant interactions with the conserved domains of RNAP to stabilize the complexes (**Figs. 1-3** and **Supplementary Figs. 13**-17).

The most amazing feature is that PhoP adopts as novel tandem dimers to specifically recognize different number of PHO boxes included in the concerning promoters (**Figs. 1, 2**, and **Supplementary Figs. 15**, 16c, 16e, 17a**, 17c, 17e**). As observed for most OmpR*/*PhoB subfamily regulators^21,32^, in the structure of 2PhoP-TAC, PhoP^II^_DBD and PhoP^I^_DBD form a head-to-tail dimer and interact with the a1 and b1 sites overlapped with the -35 element. The long helix of each PhoP_DBD inserts into the DNA major groove via an array of residues N212, E215, S216, Y217, V218, S219, Y220, R222, R223, and the C-terminal anti-parallel β sheets flap into the adjacent downstream minor groove by virtue of the residues T235, R237, G238, V239, and Y241 (**Fig. 2b)**. Interactions between the flapped residues S175, P176, T177 and the DNA backbone phosphates also assist these protein-DNA interactions. Moreover, the dimer is also stabilized by the polar and hydrophobic intersubunit interfaces constituted by residues E161, T162, H163, E164, P176 from PhoP^I^_DBD and residues N188, G190, T191, V192, R244, G238, V239 from PhoP^II^_DBD (**Fig. 2c** and **2d)**. In good agreement with this, alanine substitutions of the involved residues (E161, E164, P176, and R237) repressed PhoP-dependent transcription activities (**Fig. 2e**), supporting the importance of these residues in dimerization and DNA engagement of PhoP. Further comparisons between the PhoP-TACs and the *Mtb* RPo also unveil a remarkable difference—insertion of PhoP^I^_DBD and PhoP^II^_DBD into the adjacent major grooves nearby the -35 element significantly raises the DNA strands, replacing the canonical specific contacts between σ^A^R4 and the -35 element in most bacterial RPo (**Fig. 2f** and **2g**). Possibly due to this, the PhoP proteins confer the ability to drive RNAP to initiate transcription of the PhoP target promoters instead of host-keeping genes.

**Fig. 2.**
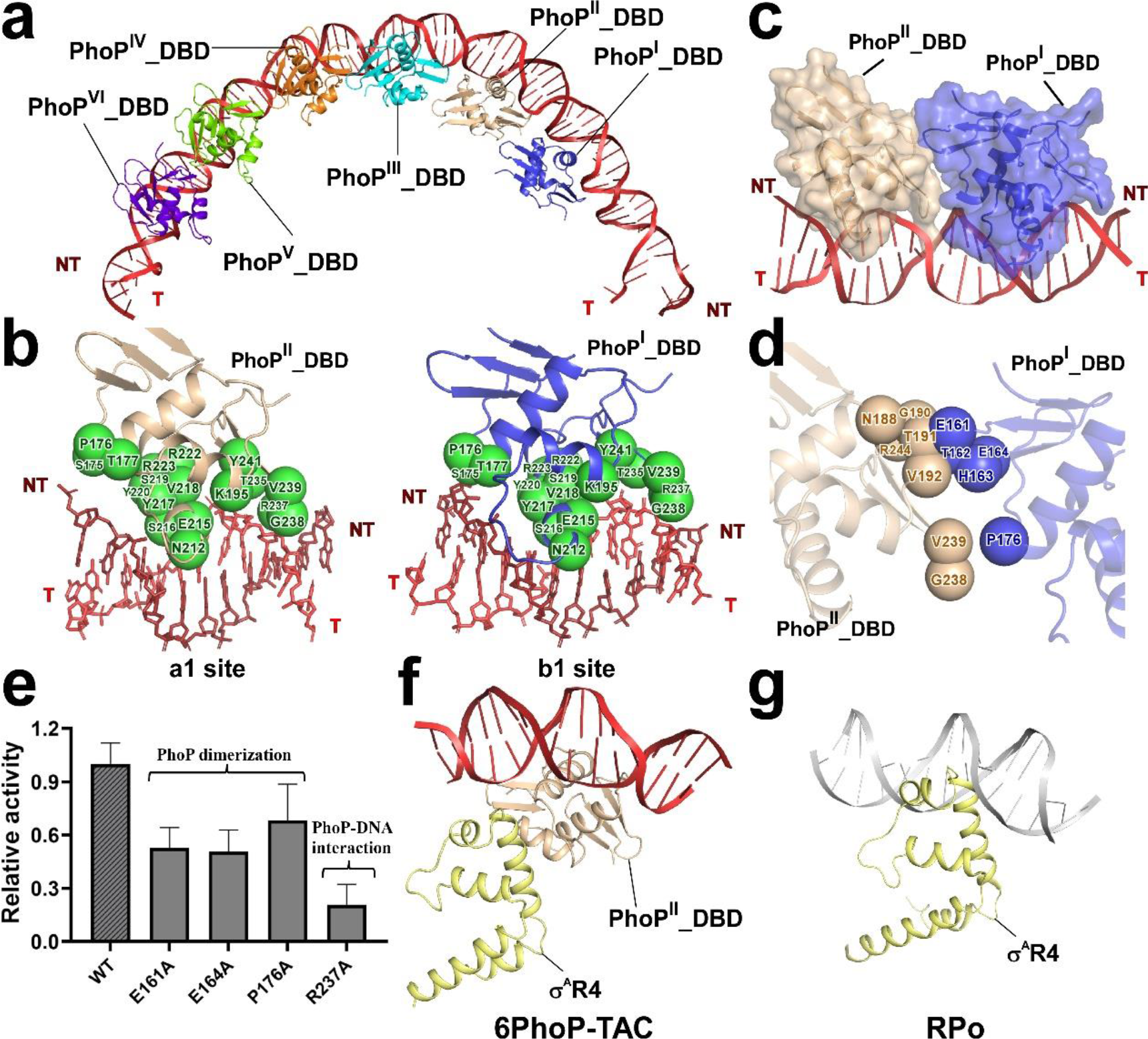
Tandem PhoP hexamer engages promoter DNA in *Mtb* 6PhoP-TAC. **a,** Relative locations of *Mtb* PhoP^I^_DBD, PhoP^II^_DBD, PhoP^III^_DBD, PhoP^IV^_DBD, PhoP^V^_DBD, PhoP^VI^_DBD located at the upstream double-stranded DNA. **b,** Detailed interactions between *Mtb* PhoP^I^_DBD, PhoP^II^_DBD, and their corresponding PhoP binding sites. The key residues involved are shown as green spheres. (**c**, **d**) The relative locations and detailed interactions between PhoP^I^_DBD and PhoP^II^_DBD bound to the promoter DNA. The key residues involved in PhoP^I^_DBD and PhoP^II^_DBD are shown as wheat and blue spheres, respectively. **e,** Substitutions of PhoP residues involved in promoter engagement and dimeric interface reduce *in vitro* transcription activities. **f,** Relative locations of PhoP^II^_DBD, σ^A^R4, and the upstream double-stranded DNA in *Mtb* 6PhoP-TAC. **g,** Relative locations of σ^A^R4 and the upstream typical -35 element DNA in *Mtb* RPo (PDB ID: 6VVY). Colors are shown as in **Fig. 1c**.

By virtue of similar protein-DNA and dimeric interactions, a tandem PhoP tetramer engages promoter DNA and drives RNAP to form 4PhoP-TAC (**Figs. 1b, 2** and **Supplementary Figs. 16**c**, 16e**), which highly resembles our recently reported cryo-EM structure of *Mtb* GlnR-TAC (PDB ID: 8HIH)^7^. However, the upstream DNA in 4PhoP-TAC is not pulled by a downward αCTD, leading the DNA strand about 2 nm (∼a helix diameter) further from RNAP core enzyme than that in GlnR-TAC (**Supplementary Figs. 19**a). Even though, the PhoP-REC domain surrounding the N-terminal domain of RNAP α subunits (αNTDs) bridges RNAP and the distal PhoP molecules, which tends to compensate and stabilize the structure of 4PhoP-TAC. While most strikingly, a novel tandem PhoP hexamer makes specific contacts with three typical PHO boxes (including a1, b1, a2, b2, a3, and b3 sites) in the structure of 6PhoP-TAC, which coordinately induces a large DNA distortion of the upstream DNA by combination with two PhoP-REC domains adjacent to RNAP αNTDs (**Figs. 1c, 2** and **Supplementary Figs. 17**a, 17c, 17e). Though the density quality of PhoP-RECs is not so high, the buried surface areas between PhoP-REC domains and RNAP αNTD are ∼350 Å^2^ and ∼120 Å^2^, respectively. This indicates the PhoP-RECs oligomerize and cooperatively bridge the PhoP_DBDs and the main body of RNAP, which further stabilizes 6PhoP-TAC and enhances transcription of the target genes.

Except for the aforementioned interactions, all of the PhoP-TACs are also stabilized by substantial protein-protein interactions between PhoP and RNAP conserved domains consisting of β flap, σ^A^R4, and αCTD (**Fig. 3** and **Supplementary Figs. 16**b, 16d, 17b, 17d), suggesting their significance and conservation in stabilizing PhoP-TACs. A cluster of residues (I187, N188, G190, T191, V192, K197, and H201) from PhoP ^I^_DBD make polar and hydrophobic contacts with the residues P808, G810, K763, L764, G765, G810, E811, and E831 from the loop of RNAP β flap, respectively (**Fig. 3a**). Meanwhile, residues Y205, D206, and F207 from PhoP^I^_DBD make electrostatic interactions with residues Q485, P486, R487, T488, and R478 from σ^A^R4, while the residues D200, H201, D206, F207, G208, G209, D210, P196, L199 from PhoP^II^_DBD also interact with residues E501, R502, R504, Q505, S508, K509, and S512 from the conserved C-terminal helix of σ^A^R4 (**Fig. 3b**). These share similarities with the interactions occurred in GlnR-TAC. Nevertheless, the αCTD is only observed in 6PhoP-TAC rather than in 2PhoP-TAC and 4PhoP-TAC, reflecting a more stable PhoP-TAC assembled (**Fig. 3c)**. Consistently, site-directed mutagenesis analysis of the involved residues (K197A, H201A, D206A, and F207A) largely reduces PhoP-dependent transcription activities, especially for D206A, and F207A, suggesting the crucial necessity of these interfaces in promoting transcription activity of PhoP (**Fig. 3d)**.

**Fig. 3.**
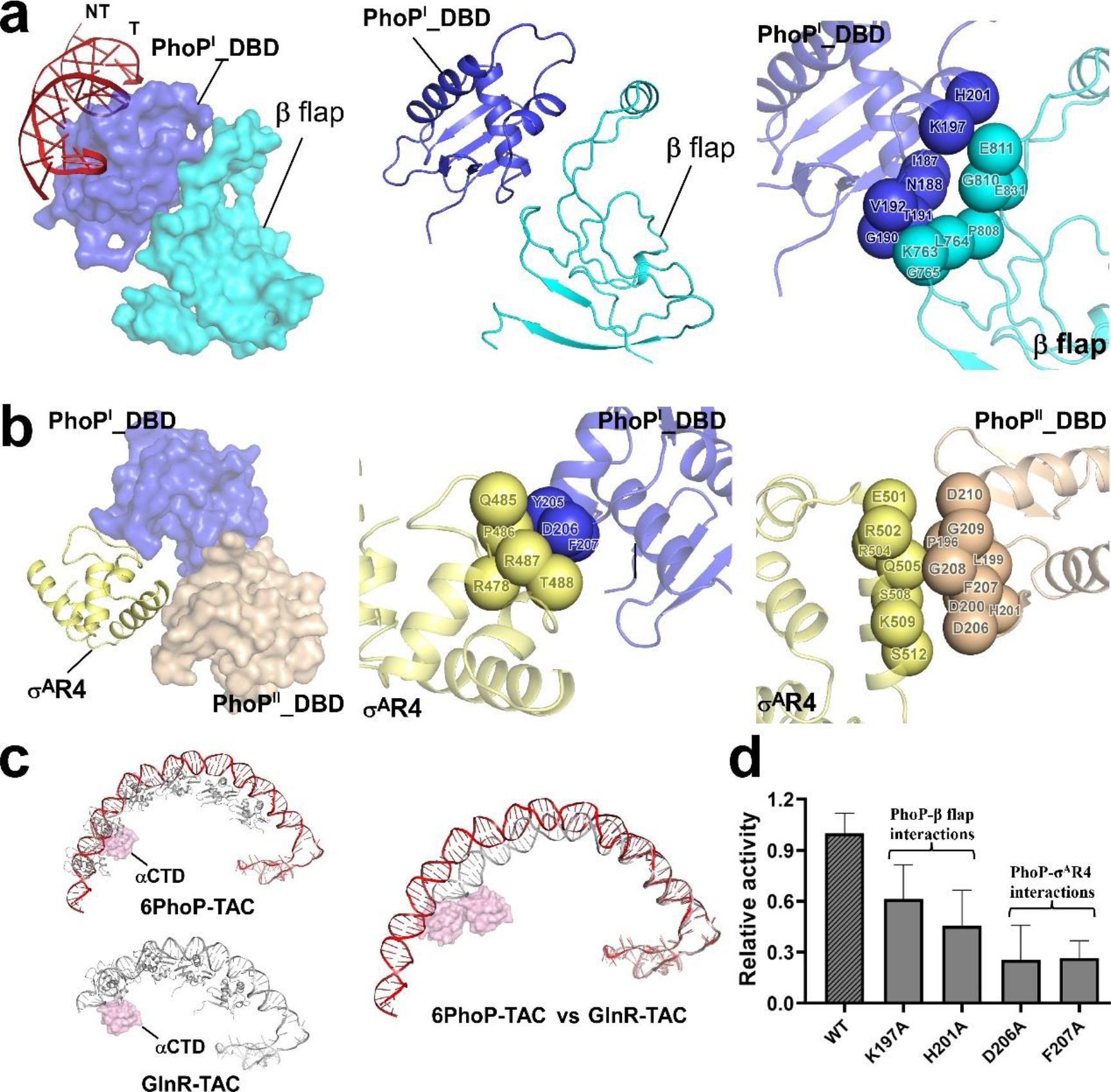
The critical protein–protein interactions between *Mtb* PhoP and the conserved β flap and σ^A^R4 domains of RNAP. **a,** Relative locations of *Mtb* PhoP^I^_DBD and RNAP β flap (Left and Middle). PhoP^I^_DBD and RNAP β flap are shown in surface style (Left). Residues involved between PhoP^I^_DBD and RNAP β flap are shown as slate (PhoP^I^_DBD) and cyan (RNAP β flap) spheres (Right). **b,** Relative locations of *Mtb* PhoP_DBDs and RNAP σ^A^R4 (Left and Middle). PhoP^I^_DBD, PhoP^II^_DBD and RNAP σ^A^R4 are shown in surface (Left). Residues involved in interactions between PhoP_DBDs and RNAP σ^A^R4 are shown as blue (PhoP^I^_DBD), wheat (PhoP^II^_DBD) and yellow (RNAP σ^A^R4) spheres (Middle and Right). RNAP β flap is colored in cyan, and RNAP σ^A^R4 is colored in yellow. **c,** Conformational comparisons of αCTD and DNA from 6PhoP-TAC and GlnR-TAC. gray, PhoP or GlnR; pink, αCTD; red, DNA from 6PhoP-TAC; gray, DNA from GlnR-TAC. **d,** Substitutions of PhoP residues involved in PhoP-β flap, PhoP- σ^A^R4 interfaces compromise *in vitro* transcription activities. The other colors are shown as in **Fig. 1**.

### Cryo-EM analysis of PhoP-TRC reveals a novel mode of bacterial transcription repression

To uncover the mysterious cross-talk veil of the regulation mechanism of the essential phosphate-nitrogen metabolism controlled by the global transcription factors PhoP and GlnR, we resolve the cryo-EM structure of PhoP-TRC and sheds new lights on a novel mode of bacterial transcription repression (**Fig. 4** and **Supplementary Figs. 7**, 12, 18, 19b).

**Fig. 4.**
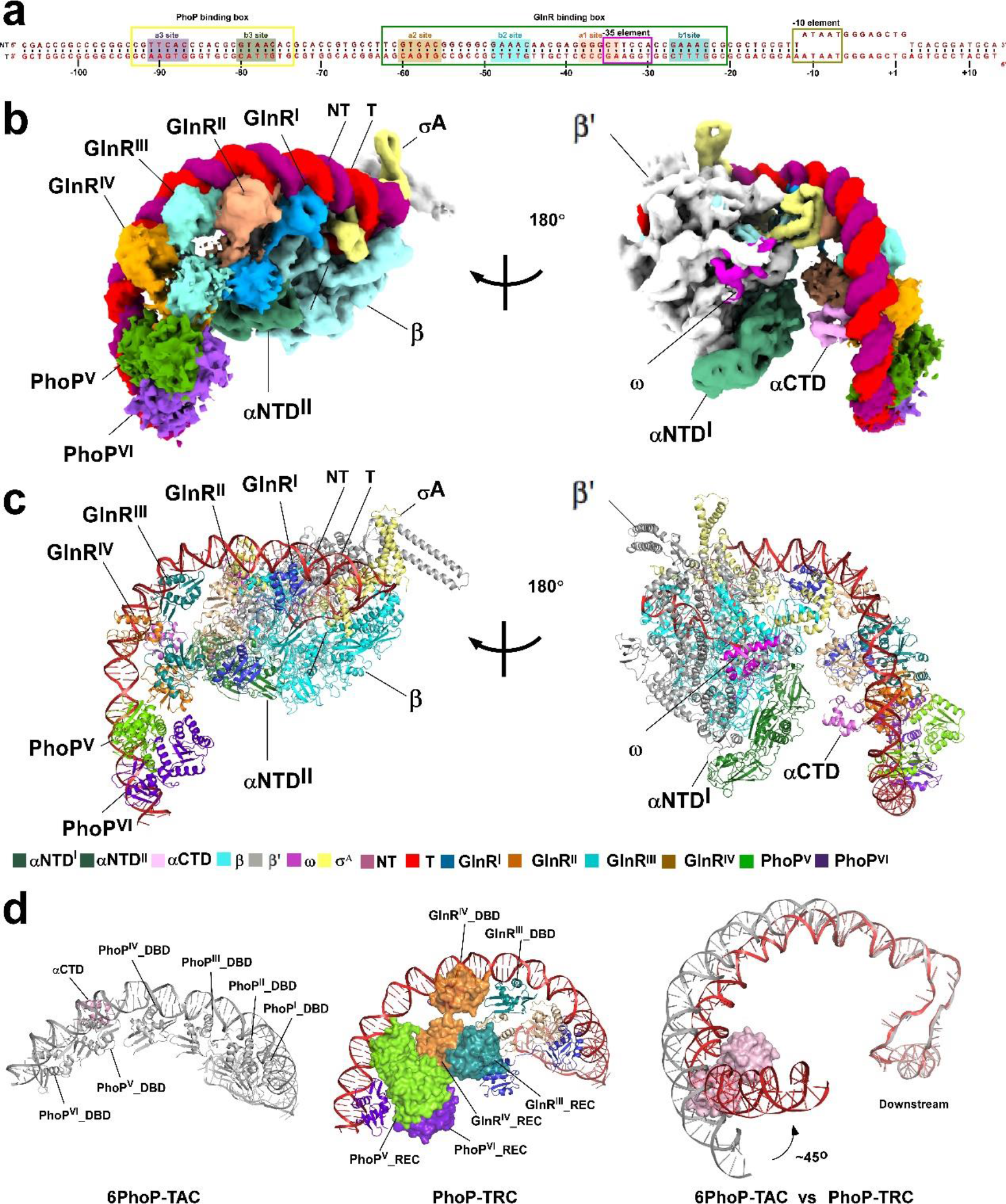
The overall structure of *Mtb* PhoP-TRC. **a,** DNA scaffold used in structure determination of *Mtb* PhoP-TRC. GlnR binding box and PhoP binding box are framed in green and yellow color, respectively. The a1 site, b1 site, a2 site, b2 site, a3 site, b3 site are shaded in light orange, dark cyan, brown, cyan, purple, and dark green, respectively. **(b, c)** Two views of the cryo-EM density map **(b)** and structure model **(c)** of *Mtb* PhoP-TRC. The EM density map is colored as indicated in the color key. **d,** The promoter DNA bends toward the main body of RNAP in PhoP-TRC compared with the structure of *Mtb* 6PhoP- TAC. The other colors are shown as in **Fig. 1**.

In PhoP-TRC, four GlnR molecules collaboratively engage the a1, b1, a2, and b2 sites of the PhoP-GlnR co-regulated promoter *amtB*, and simultaneously make various types of protein-protein interactions with the conserved domains (β flap, σ^A^R4, αCTD, and αNTD) of RNAP, highly resembling those interactions observed in the structure of *Mtb* GlnR-TAC (**Fig. 4b, 4c** and **Supplementary Figs. 18**, 19b). While it is quite remarkable that one more PhoP dimer (PhoP^V^ and PhoP^VI^) interacts with the a3 and b3 sites via the same sets of PhoP-DNA interactions as those in PhoP-TACs (**Supplementary Figs. 18**h). Nevertheless, the spacer between the b3 site and a2 site is 15 bp in length, which is much longer than the other 6 bp- spacers downstream or between PhoP^V^_DBD and PhoP^VI^_DBD in 6PhoP-TAC (**Fig. 4d**, left panel). This unique longer spacer allows for PhoP^V^_DBD to interact with the GlnR^IV^_REC, and the dimeric PhoP_RECs to make contacts with the adjacent GlnR_RECs, presenting the overlapped surface areas of ∼95 Å^2^ and ∼168 Å^2^, respectively (**Fig. 4d**, middle panel). These special protein-DNA and protein-protein interactions from the PhoP dimer drag the upstream portion of promoter DNA distorting toward RNAP core enzyme with ∼45°, which is most prone to be a negative force to drag forward movement of RNAP (**Fig. 4d** right panel and **Supplementary Figs. 19**b, 19c). Therefore, it urges us to speculate that PhoP probably acts in a “dragging model” to repress transcription initiation. To find out whether the longer spacer is essential for PhoP-TRC, we shortened the spacer by half a helix (del 5 bp) or one helix (del 10 bp), and assessed their corresponding effects on formation of the PhoP-TRC via EMSA. Consistent with our hypothesis, it yielded fewer migration bands of PhoP-TRC with the del 5 bp promoter than with the del 10 bp promoter, and the wild type WT promoter (**Supplementary Figs. 2**h), revealing the dependence of such a longer spacer on PhoP-dependent transcription repression.

## Discussion

During evolution, the pathogenic *Mtb* has seized intricate transcriptional regulation network to swiftly govern differential expression of stress genes, which is critical for the resultant bacterial colonization and infection^4,6^. This complex stress adaptation mechanism leads to diverse gene variability which is generally termed as transcriptional plasticity. Recently, transcriptomic analysis of *Mtb* genome under a multitude of environmental pressures points out that the profile of significant transcriptional plasticity closely correlates with transcription regulators, possible promoter features and relevant regulatory mechanisms^33^. However, the intricate molecular mechanisms underlying such stress adaptation are still obscure. In the present work, we succeed in determining three cryo-EM structures of *Mtb* PhoP-TACs, one cryo-EM structure of *Mtb* RPo, and one cryo-EM structure of *Mtb* PhoP-TRC (**Figs. 1**, **4** and **Supplementary Figs. 13**), and obtain a deep understanding on the structural basis of the long-standing questions based on the structural and biochemical findings.

Transcriptomic and genetic identification reveal that most bacterial stress promoters contain suboptimal -35 elements, which cannot be recognized with high affinity by the predominant σ^A^. Even though, this can be mainly compensated by abundant transcription activators. By comparison with *Mtb* RPo, the structures of PhoP-TACs shed new structural insights into PhoP-dependent transcription activation: phosphorylation of PhoP is required to trigger potential conformation changes, enhancing DNA binding and promoting formation and stabilization of active PhoP-TACs, which is also favored by additional protein-protein interactions between the REC domains of PhoP. Though σ^A^R4 is unable to make direct interactions with the suboptimal -35 element as those observed in most bacterial RPo, the downstream PhoP dimer takes this responsibility through specifically recognizing the PhoP target promoters and simultaneously interacting with the conserved domains of RNAP (β flap, σ^A^R4, αCTD, and αNTD) (**Figs. 1** and **2**). Notably, the PhoP proteins also distinctively adopt in tandem dimer/tetramer/hexamer oligomerization state to strengthen the specific replacements at promoters containing different number of PHO boxes (**Fig. 5a**), which are quite different from the reported monomeric or dimeric transcription activators ^26,30,34,35,36,37,38^. Though 2PhoP-TAC and 4PhoP-TAC share similarities with the PmrA-TAC and GlnR-TAC in protein-DNA and protein-protein interactions, the 6PhoP-TAC represents a novel mode of bacterial transcription activation, in which the activators bind both upstream and downstream of the suboptimal -35 element as a tandem hexamer. Since similar observations are also observed in the GlnR-TAC and PmrA-TAC^7,19^, it is conceivable that this unique activator- replacement manner may represent a general transcription activation mode for the OmpR*/*PhoB family regulators in actinobacteria. As also coincides with and enriches the “Type IV σ^70^ family- dependent transcription activation mechanism’’ reviewed recently by Liu, *et al*^26^. As to the physiological significance of these diversified regulatory modes of the tandem PhoP dimers, we infer that these architectures are prone to exhibit more superiority to differentially evoke transcription initiation of more cellular PhoP target regulons, respond to the surrounding environmental stresses with higher sensitivity and efficiency, and thus greatly improve bacterial survival under adversity.

**Fig. 5.**
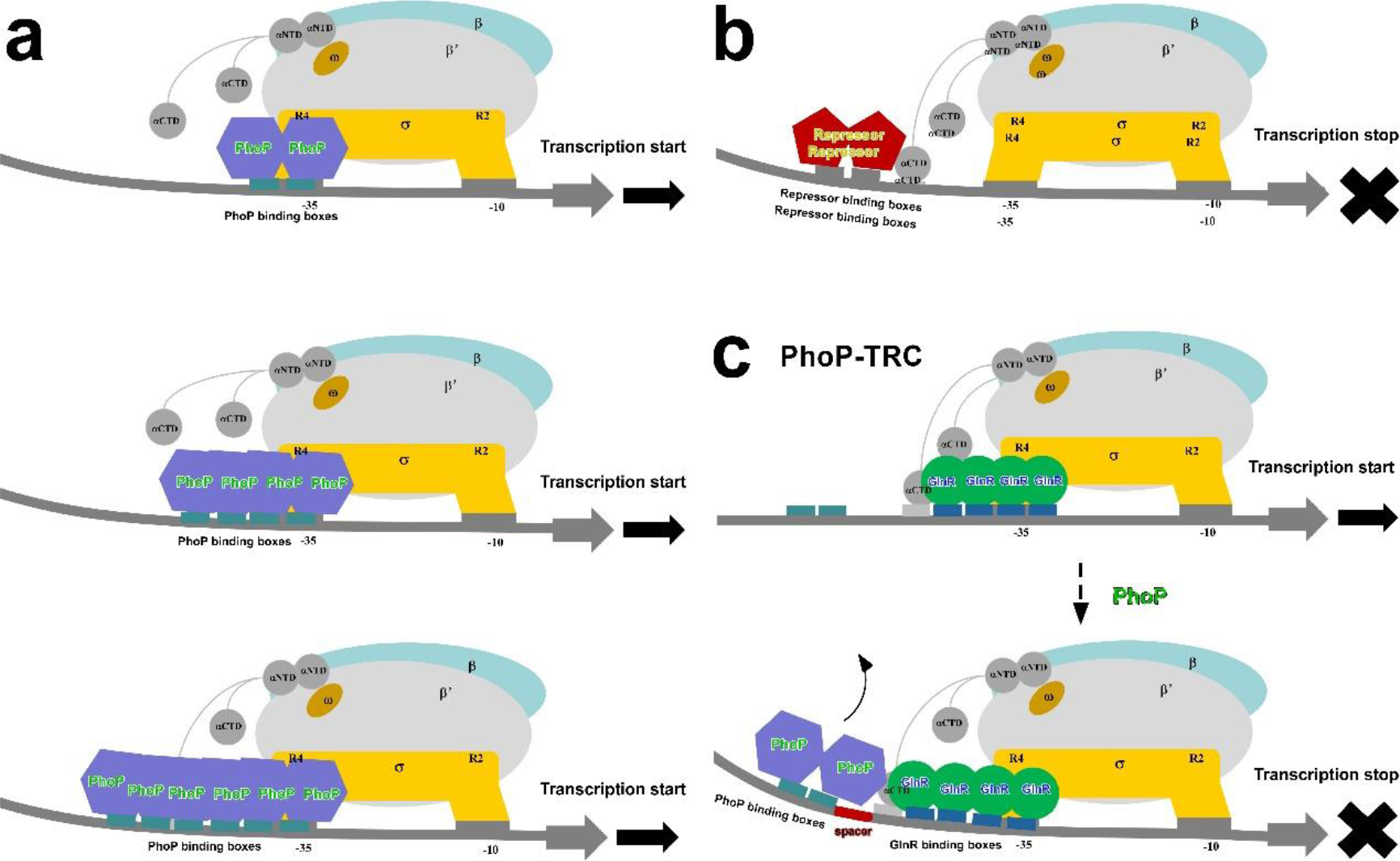
Proposed working model for PhoP-dependent transcription regulation. **a,** Three types of PhoP-dependent transcription activation in which PhoP adopts as tandem dimer, tetramer, and hexamer, respectively. **b,** Classic transcription repression model by occupying RNAP binding sites at a promoter. **c,** A proposed “dragging model” for PhoP-dependent transcription repression. In the absence of the upstream PhoP dimer, four GlnR molecules cooperatively activate transcription of the target promoter. When PhoP dimer specially interacts with the a3 and b3 sites, it may distort the upstream DNA and drag the GlnR molecules through extensive protein-DNA and protein-protein interactions, which may impede formation and stabilization of a stable TAC, exert negative force to forward movement of RNAP, and thereby repress and transcription initiation of the downstream genes. Repressor and spacer are shown as five-pointed star and rectangle, and colored in red.

Different from the reported transcription repressors those occupying promoter consensus binding elements to block RNAP (**Fig. 5b**)^39,40^, cryo-EM structure analysis of the PhoP-TRC highlights the molecular determinants of PhoP-dependent transcription repression on the cross-talked nitrogen metabolism promoter: despite of the conserved PhoP-PHO box interactions, the upstream promoter spacer length between PhoP and GlnR, along with the PhoP-GlnR and PhoP-RNAP αNTD interactions collectively contribute to formation and stabilization of the PhoP-TRC. Thus, it is tempting to speculate that PhoP may act in an utmost reasonable “dragging model” to repress transcription initiation (**Fig. 5c)**: in the absence of the upstream PhoP dimer, four GlnR molecules cooperatively activate transcription of the target promoter. When PhoP proteins appear, they dimerize and specially interact with the a3 and b3 sites, drag the upstream DNA and the GlnR molecules through extensive protein-DNA and protein-protein interactions (**Fig. 4** and **Supplementary Figs. 18**, 19b). Thereby, this framework cooperatively ensures PhoP in a novel “dragging model” to impede formation and stabilization of a stable TAC, exhibit negative force for forward movement of RNAP, and thus repress transcription initiation of the downstream genes. This well coincides with the deduction about the upstream a3-b3 site which may serve as an elaborate transcription strategy to swiftly respond to cellular nutritional starvation. Such a finely tuned coordination mechanism highlights the structural basis for balancing phosphate–nitrogen utilization and interplay, which ensures overall ‘metabolic homeostasis’ to sustain bacterial survival.

As verified by both *in vivo* and *in vitro* investigations, the conserved mutation S219L (Ser219 replaced by Leu) of the global virulence regulator PhoP in the avirulent *Mtb* H37Ra strain impairs synthesis of several virulent cell wall lipids and alters its morphology, which are proved to contribute to its virulence attenuation compared with the virulent *Mtb* H37Rv strain^41, 42, 43^. In good agreement with the physiological importance, the residue S219 from the long DNA binding helix of *Mtb* PhoP makes specific interactions with the bases of the template strand DNA in the structures of both PhoP-TACs and PhoP-TRC (**Fig. 2b** and **Supplementary Figs. 16**c, 16e, 17a, 17c**, 17e, 18h**), confirming the crucial necessity of the highly conserved residue, and also revealing the functional states of these PhoP-dependent transcription regulation complexes.

Taken together, the structural and biochemical data not only reveal the critical significance of the genetic promoter characteristics (number of PHO boxes, consensus nucleotides, and intervening spacer lengths), PhoP oligomerization, the conserved PhoP-PHO box interactions, and extensive protein-protein interactions between PhoP and PhoP/RNAP/GlnR in determining PhoP as an activator or a repressor to deftly tune transcription initiation of stress genes, but also expand our understanding on the intricate transcriptional regulation network governing persistence of the pathogenic *Mtb* to host challenging stresses. Moreover, these various modes of PhoP-dependent transcription regulation provide new structural evidences for the global OmpR/PhoB family regulator in fostering high sensitivity and variability of bacterial transcriptional plasticity. Since the global virulence factor PhoP is highly conserved among *Mtb* and the other leading pathogens, our findings may guide further development of innovative strategies or interventions targeting PhoP to combat the resulting bacterial infections.

## Methods

### Plasmids and DNA

In order to construct the expression plasmid of pET28a-*phop*, the *Mtb* PhoP encoding *gene* fused with an N-terminal His×6 tag under the control of T7 promoter was synthesized into pET28a by Sangon Biotech, Inc. The recombinant plasmids containing the site-directed *Mtb* PhoP mutants were constructed according to the manual of the QuikChange Site-Directed Mutagenesis Kit, Agilent, Inc. To compare transcription activities of PhoP on promoters containing different number of PhoP binding sites (2 sites, 4 sites, 6 sites), the phosphate metabolism promoter *phoU* encompassing 2 sites (a1-b1), 4 sites (a2-b2 and a1-b1), and 6 sites (a3-b3, a2-b2, and a1-b1), followed by an RNA aptamer (Mango III) coding sequence, were amplified and purified using the QIAquick PCR Purification Kit (Qiagen, Inc.). These DNA fragments were designated as *phoU2*, *phoU4*, and *phoU6*, respectively. Nucleic-acid scaffolds used to assemble *Mtb* RPo, *Mtb* PhoP-TACs, and *Mtb* PhoP-TRC were annealed from the synthetic template-strand DNA (T) and non-template-strand DNA (NT) via an annealing procedure (95 °C, 5 min followed by 2 °C-step cooling to 25 °C) in the annealing buffer (20 mM Tris-HCl, pH 8.0, 200 mM NaCl). Primers used in this study are shown in **Supplementary Table S1**.

### *Mtb* PhoP

Plasmids of pET28a-*phop* or pET28a-*phop* derivative were first transformed into BL21(DE3) (Invitrogen, Inc.). Then, single colonies of the positive transformants were inoculated and amplified with 5 L LB broth supplemented with 100 μg/mL kanamycin at 37 °C with shaking. When OD600 of the cultures reached about 0.8, PhoP expression was induced by adding 0.5 mM IPTG, and cultures were incubated overnight at 20 °C. After being harvested by centrifugation (5,000 rpm; 15 min, 4 °C), the cell pellets were resuspended in 100 mL buffer A (20 mM Tris–HCl, pH 7.9, 300mM NaCl, and 5% glycerol) and lysed using an ATS AH-10013 cell disrupter (ATS, Inc.). After centrifugation at 12,000 rpm for 30 min at 4 °C, the supernatant was loaded onto a 5-mL column of Ni-NTA agarose (Qiagen, Inc.) pre-equilibrated with buffer A. The column was washed with 10 mL buffer A containing 40 mM imidazole and eluted with 30 mL buffer A consisting of 200 mM imidazole. Then, the elutes were concentrated and purified by a 120-mL HiLoad 16/600 Superdex 75 column (GE Healthcare, Inc.) with buffer B (20 mM Tris–HCl, pH 7.9, 300 mM NaCl, 5 mM MgCl2, and 1mM DTT). After verification by SDS-PAGE, the target elutes containing PhoP were pooled and stored at –80 °C. Yield was ∼1.5 mg/L, and purity was > 90%. To enhance the relative DNA binding affinity, the purified PhoP proteins were phosphorylated by incubating with 50 mM AcP at 30 ℃ for 90 min. After centrifugation at 12,000 rpm for 30 min at 4 °C, the sample was loaded onto a 25- mL Hitrap desalting column (Cytiva, Inc.) and eluted with the same buffer B to remove extra AcP. Finally, the phosphorylated PhoP proteins were pooled and stored as described above. Similarly, the phosphorylated PhoP derivatives were prepared. To be explicit, the following phosphorylated PhoP and phosphorylated PhoP derivatives were designated as PhoP and PhoP derivatives, respectively.

### *Mtb* RNAP

*Mtb* RNAP holoenzyme including σ^A^ was induced from cultures of *E. coli* strain BL21(DE3) (Invitrogen, Inc.) co-transformed with plasmids of pACYC Duet-*rpoA*-*sigA*, pCDF-*rpoZ*, and pET Duet-*rpoB*-*rpoC*, and purified as described previously ^7^.

### Assembly of PhoP-TAC and PhoP-TRC

The DNA scaffolds *pstS* (containing 2 PhoP binding sites), *phoU4* (containing 4 PhoP binding sites), *phoU6* (containing 6 PhoP binding sites) were used to assemble 2PhoP-TAC, 4PhoP-TAC, and 6PhoP-TAC, respectively. Likewise, the DNA scaffold *amtB* that consists of 2 PhoP binding sites and 4 GlnR binding sites were chosen to assemble the repression complex PhoP-TRC. Firstly, the PhoP-TACs were assembled by incubating *Mtb* RNAP, DNA scaffold, and PhoP in a molar ratio of 1: 1: 8 at 4 °C overnight. The sample was then applied onto a 120- mL HiLoad 16/600 Superdex 200 column (GE Healthcare, Inc.) and eluted with buffer C (20 mM Tris–HCl, pH 7.9, 75 mM NaCl, 5 mM MgCl2, and 1mM DTT). Once being verified by SDS-PAGE, the fractions containing each component of the assembled PhoP-TAC were concentrated using Amicon Ultra centrifugal filters (10 kDa MWCO, Merck Millipore, Inc.). In addition, the PhoP-GlnR complex was assembled by incubating *Mtb* RNAP, *amtB* scaffold, PhoP and GlnR in a molar ratio of 1:1:8:16 at 4 °C overnight, and subsequently purified by a 120-mL HiLoad 16/600 Superdex 200 column and assessed by SDS-PAGE. Oligonucleotides synthesized to prepare the DNA scaffolds are shown in **Supplementary Table S1**.

### Cryo-EM grid preparation

Firstly, the Quantifoil grids (R1.2/1.3 Cu400 mesh; Quantifoil, Inc.) were glow-discharged for 120 s at 25 mA. Then, 3 μL of each purified *Mtb* PhoP-dependent transcription regulation complex was applied onto the grids after incubation with 8 mM CHAPSO (Hampton Research Inc.) for 1 min at 25 ℃. Subsequently, the grids loaded with samples were blotted with Vitrobot Mark IV (FEI), immediately plunge-frozen into liquid ethane with 95 % chamber humidity at 10 ℃. Finally, the grids screened with moderate density and uniform distribution of single particles will be chosen for large amounts of cryo-EM data collection.

### Cryo-EM data acquisition and processing

Cryo-EM data of each *Mtb* PhoP-dependent transcription regulation complex was acquired with the same set of parameters on a 300 kV Titan Krios (FEI, Inc.) equipped with a K3 Summit direct electron detector, and sequentially processed with the relevant cryo-EM data analysis software (**Table 1**, and **Supplementary Figs. 3**-7). Different number of images were recorded with EPU in counting mode with a pixel size of 1.2 Å, a dose rate of 10 e/pixel/s, and an electron exposure dose of 50 e/Å^2^. Movies were recorded for 8.38 s with the defocus range varied from -2.0 μm to -1.0 μm. Subframes of individual movies were aligned using MotionCor2 ^44^, and contrast-transfer-function for each summed image was estimated using CTFFIND4 ^45^. From the summed images, approximately 10,000 particles were manually picked and subjected to 2D classification in RELION ^46^. The resultant 2D classes with different orientations were further chosen, auto-picked, manually inspected, and subjected to iterative 2D classification. By removing poorly populated classes, the selected particles were subjected to 3D classification in RELION by using a map of *Mtb* RPo (PDB ID: 6VVY) (for PhoP-TACs and RPo) or *Mtb* GlnR-TAC (PDB ID: 8HIH) (for PhoP-TRC) low-pass filtered to 40 Å resolution as a reference, respectively. Then, particles from the best class which shows clear density for RNAP, DNA and PhoP/GlnR were re-processed by 3D auto-refinement, re-extracted, CTF-refined, Bayesian polishing, 3D auto-refinement and post-processing in RELION. The final mean map resolutions for each *Mtb* PhoP-dependent transcription regulation complex analyzed by Gold-standard Fourier-shell-correlation are as following: 3.40 Å of RPo, 3.20 Å of 2PhoP-TAC, 4.50 Å of 4PhoP-TAC, 3.20 Å of 6PhoP-TAC, and 3.68 Å of PhoP-TRC (**Table 1** and **Supplementary Figs. 8**-12).

### Cryo-EM model building and refinement

The model of RNAP and the downstream DNA from the cryo-EM structure of *Mtb* RPo (PDB ID: 6VVY), the co-crystal structure of PhoP and DNA were fitted into the cryo-EM density map of *Mtb* RPo and PhoP-TACs using Chimera^47^. Likewise, the model of *Mtb* RNAP, GlnR and DNA from the cryo-EM structure of *Mtb* GlnR-TAC (PDB ID: 8HIH) was fitted into the cryo-EM density map of *Mtb* PhoP-TRC. The model of the upstream nucleic acids was built manually in Coot^48^. The coordinates were further calculated and validated by real-space refinement with secondary structure restraints in Coot and Phenix^49^. Structures were analyzed with Chimera and PyMOL^50^.

### Electrophoretic mobility shift assay

Electrophoretic mobility shift assay (EMSA) was carried out in EMSA buffer (40 mM Tris–HCl, pH 7.9, 100 mM NaCl, 10 mM MgCl2, and 5 % glycerol). Components and final concentration of the reaction mixture (20 μl) are as follows: 32 μM *Mtb* PhoP (or PhoP derivatives), 16 μM *Mtb* GlnR, 0.2 μM *Mtb* RNAP, and 30 nM *amtB* DNA. RNAP was firstly incubated with DNA for 10 min at 37 °C, and then incubated with GlnR or/and PhoP for 20 min at 37 °C. Once being incubated with 0.03 mg/mL heparin for 2 min at 22 °C, the reaction mixtures were applied to 5% polyacrylamide slab gels (29:1 acrylamide/bisacrylamide), electrophoresed in 90 mM Tris–borate, pH 8.0, and 0.2 mM EDTA, and stained with 4S Red Plus Nucleic Acid Stain (Sangon Biotech, Inc.) according to the procedure of the manufacturer.

### *In vitro* transcription assay

*In vitro* transcription assays were performed in transcription buffer (40 mM Tris–HCl, pH 7.9, 50 mM NaCl, 10 mM MgCl2, and 5% glycerol) by using 96-well microplates (Corning incorporated, USA). Reaction mixtures (80 μl) contained: 0.15 μM *Mtb* RNAP, 30 nM mango- ended DNA, 32 μM *Mtb* PhoP or its derivatives, 0.1 mM NTP mix (ATP, UTP, GTP, and CTP), and 1 μM TO1-biotin. AcP were used to phosphorylate *Mtb* PhoP for 90 min at 30 ℃ and then purified as described. Firstly, *Mtb* RNAP and DNA were incubated for 10 min at 37 °C. Then, PhoP was added into the mixture and incubated for 20 min at 37 °C. Subsequently, NTP mix and TO1-biotin were added, and the mixture was incubated for another 30 min at 37 °C. Finally, fluorescence emission intensities were measured using a multimode plate reader (EnVision, PerkinElmer Inc.; excitation wavelength = 510 nm; emission wavelength = 535 nm). Relative transcription activities of PhoP derivatives were calculated using

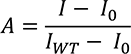

where *IWT* and *I* are the fluorescence intensities in the presence of PhoP and PhoP derivatives, respectively. *I0* is the fluorescence intensity in the absence of PhoP.

## Data Availability

The accession codes for the atomic coordinates reported in this paper are deposited in the Protein Data Bank: 8XED for RPo, 8XEC for 2PhoP-TAC, 8XEB for 4PhoP-TAC, 8X8B for 6PhoP-TAC, and 8X8D for PhoP-TRC. The accession codes for the 3D cryo-EM density maps are deposited in the Electron Microscopy Data Bank (EMDB): EMD-38287 for RPo, EMD- 38286 for 2PhoP-TAC, EMD-38285 for 4PhoP-TAC, EMD-38138 for 6PhoP-TAC, and EMD-38140 for PhoP-TRC. The PDB entries 6VVY and 8HIH are used for structure comparison in this study. Other data are available from the corresponding author upon reasonable request.

## Acknowledgements

This work was funded by the National Natural Science Foundation of China (82311530689, 32270037, 32270192, 82072240, 32000025), the Jiangsu Qinglan Project to J.S. and the Natural Science Foundation of Jiangsu Province (BK20211302, SBK2023030145), the National Key R&D Program of China (2023YFC2308200), the Open Project of Chinese Materia Medica First-Class Discipline of Nanjing University of Chinese Medicine (2020YLXK008, 2020YLXK016), the Fok Ying Tung Education Foundation, Open Funding Project of the State Key Laboratory of Bioreactor Engineering, East China University of Science and Technology to W.L.

We appreciate Shenghai Chang at the Center of Cryo-Electron Microscopy in Zhejiang University School of Medicine, Guangyi Li, Liangliang Kong, Jialin Duan, and Yun Song of the Electron Microscopy System at the National Facility for Protein Science in Shanghai (NFPS), Shanghai Advanced Research Institute, Chinese Academy of Sciences, China for providing technical support and assistance in data collection. We thank the Experiment Center for Science and Technology, Nanjing University of Chinese Medicine for experimental assistance. We thank the Core Facilities, Zhejiang University School of Medicine for technical support.

## Author Contributions

W. L., J. S., and F. Y. designed the experiments. Q. S., Z.Z. F., T.Y. L., Z.H. Y., S.M. X., F. G., and L.X. X. performed molecular cloning, protein expression and purification, enzymatic determination and cryo-EM sample assembly. A.J. W., J. S., Z.Z. F., and L.Q. X performed cryo-EM sample preparations and data acquisition. W. L. and F. Y. analyzed the cryo-EM data and determined the structures. J. S., W. L., J.P. Z. and K.D. performed formal analysis. J. S., W. L., and G.P. Z wrote the paper with help from all authors.

## Supplementary information

The online version contains supplementary material available at XXX.

---

**Supplementary Fig. 1.**
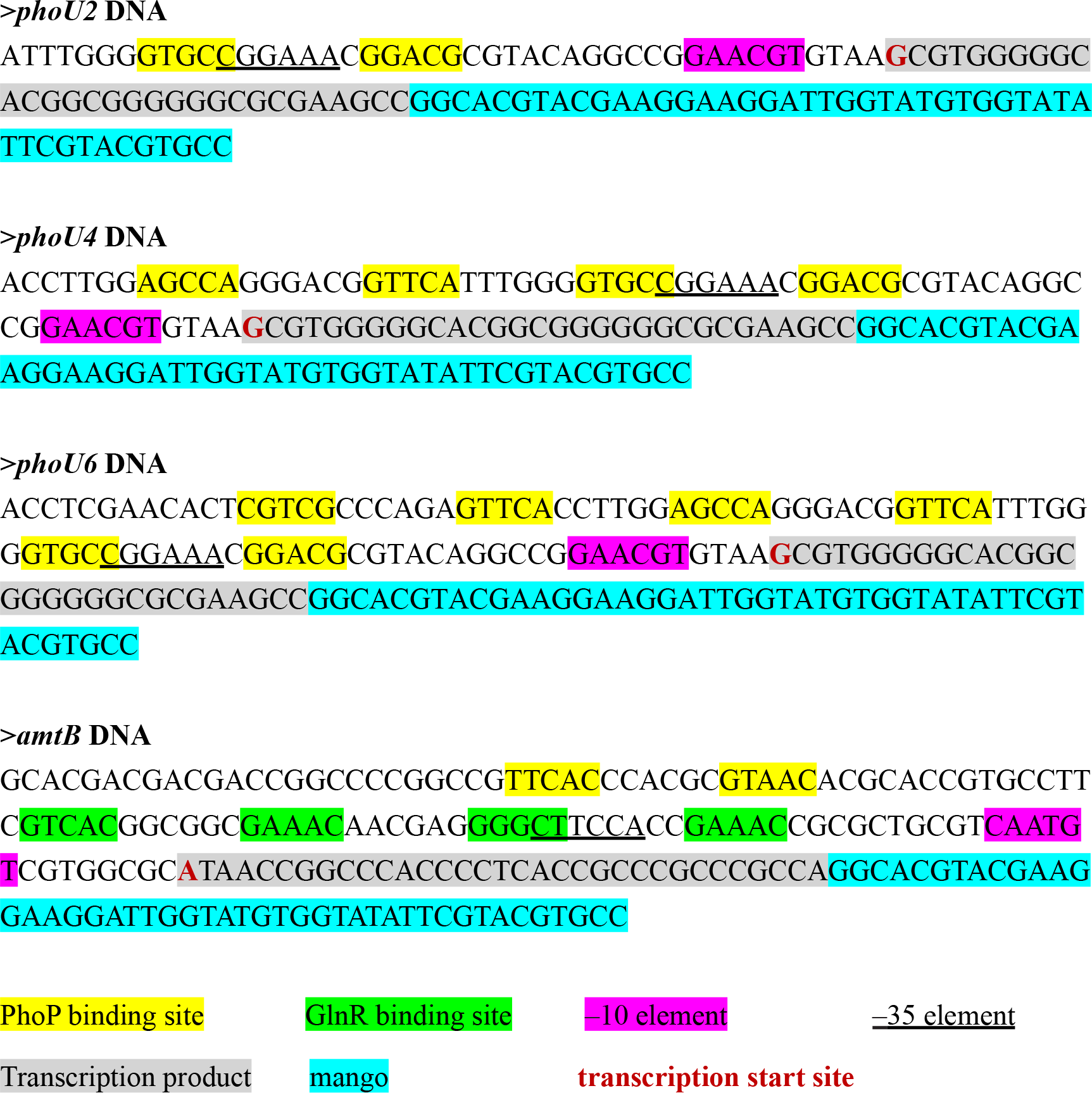
The promoter sequences used in reconstitution of PhoP-dependent transcription regulation complexes. The PhoP binding sites, GlnR binding sites, –10 element, transcription product, *mango* sequence are highlighted in green, yellow, purple, grey, and cyan, respectively. The –35 element is underlined. The transcription start site is shown in red and bold font.

**Supplementary Fig. 2.**
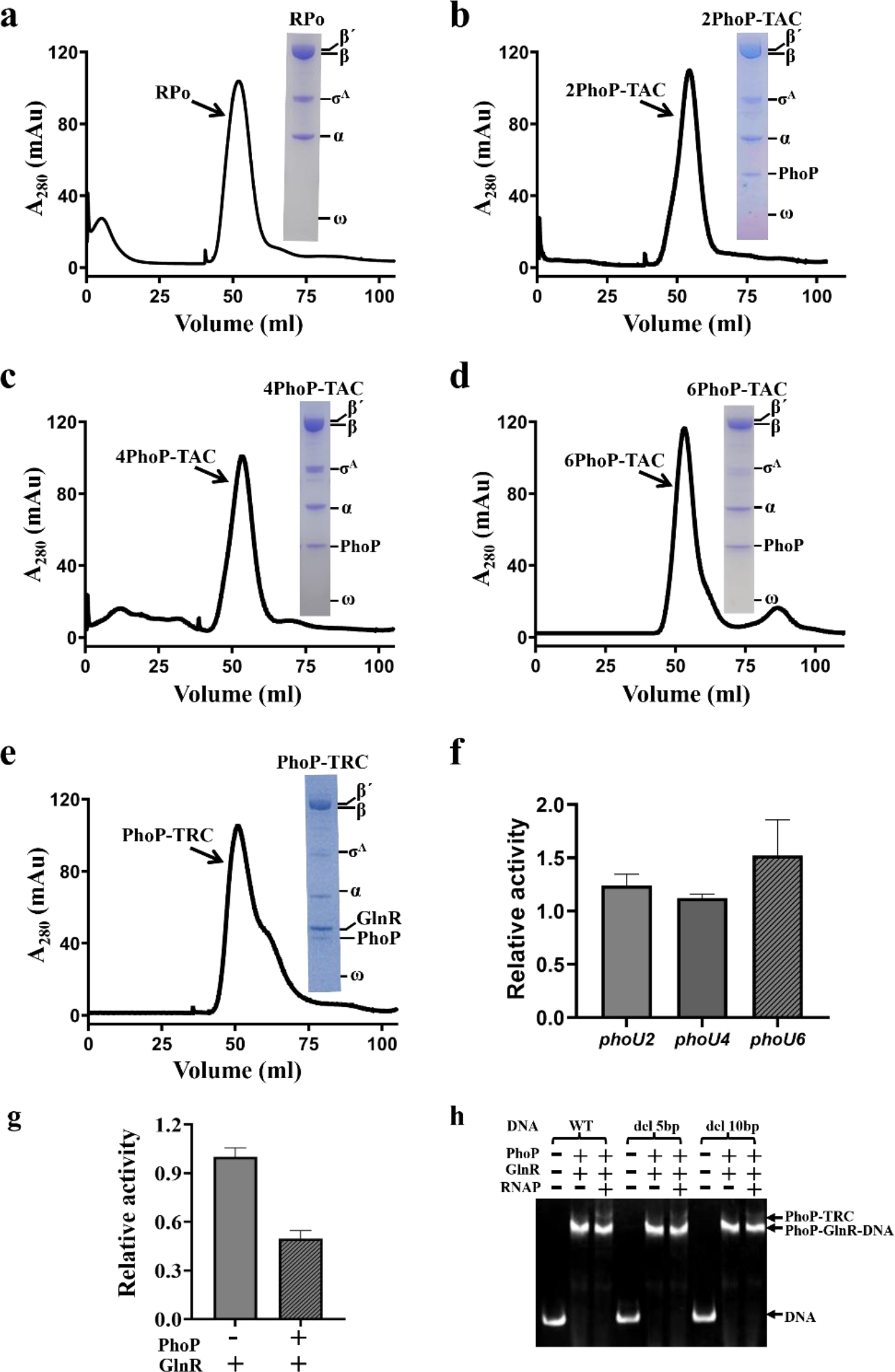
Purification and verification of *Mtb* PhoP-TACs and PhoP-TRC. a-e, Gel filtration maps and SDS-PAGE analysis of *Mtb (Mtb)* RPo, 2PhoP-TAC, 4PhoP-TAC, 6PhoP-TAC, and PhoP-TRC, respectively. **f, g,** Relative transcription activity of *Mtb* PhoP and RNAP determined by *in vitro* transcription assay on promoters containing different number of PhoP binding sites (**f**) and on the *Mtb* PhoP-GlnR co-regulated promoter (**g**). Data for *in vitro* transcription assays are means of 3 technical replicates. Error bars represent mean ± SEM of n = 3 experiments. **h,** EMSA analysis of *Mtb* PhoP-TRC formation on promoter mutants with different spacer length. Bands of PhoP-TRC, PhoP-GlnR-DNA, and free DNA are indicated by arrows on the right, respectively. Reaction conditions are described in detail in *Methods*.

**Supplementary Fig. 3.**
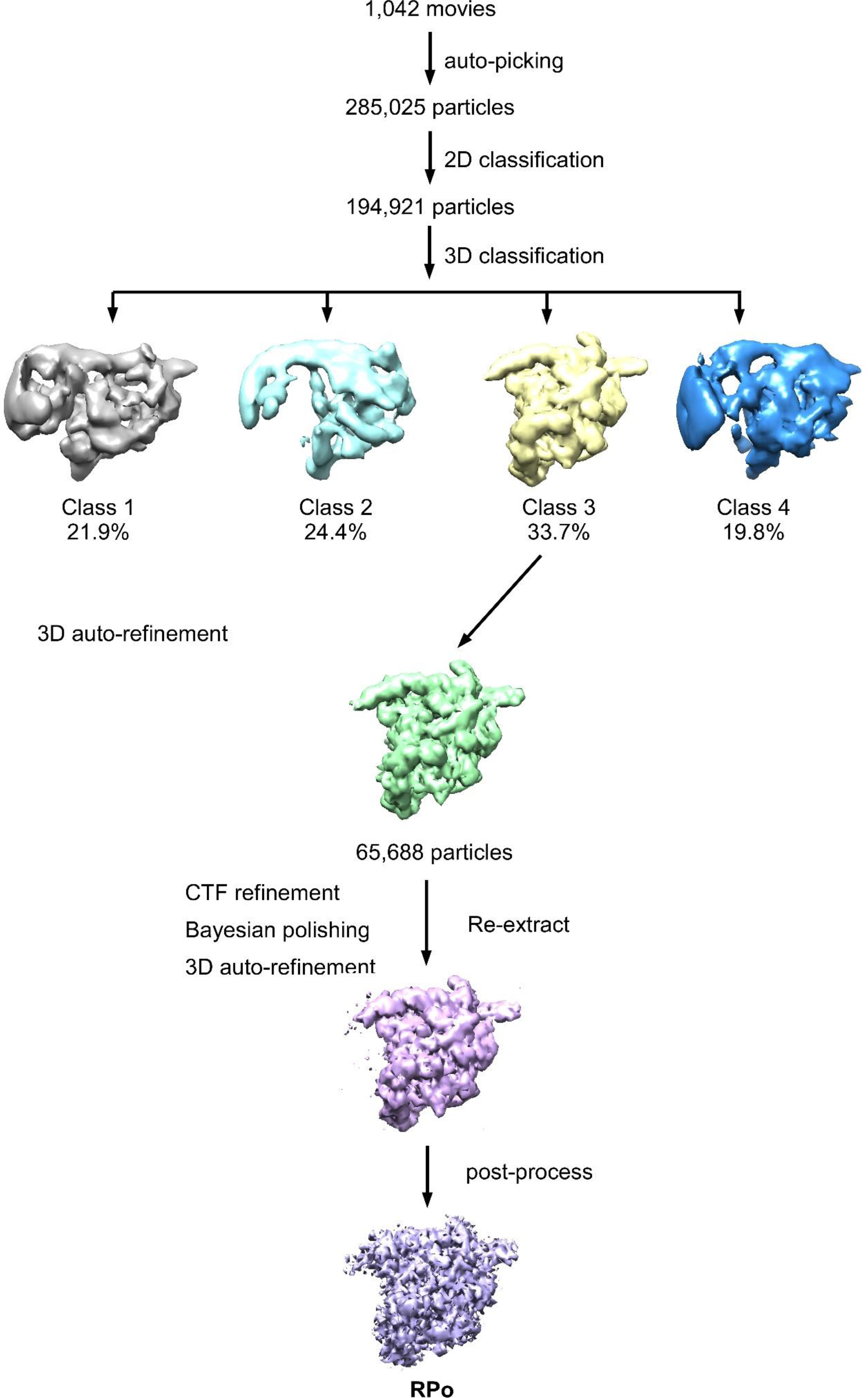
Data processing pipeline for *Mtb* RPo.

**Supplementary Fig. 4.**
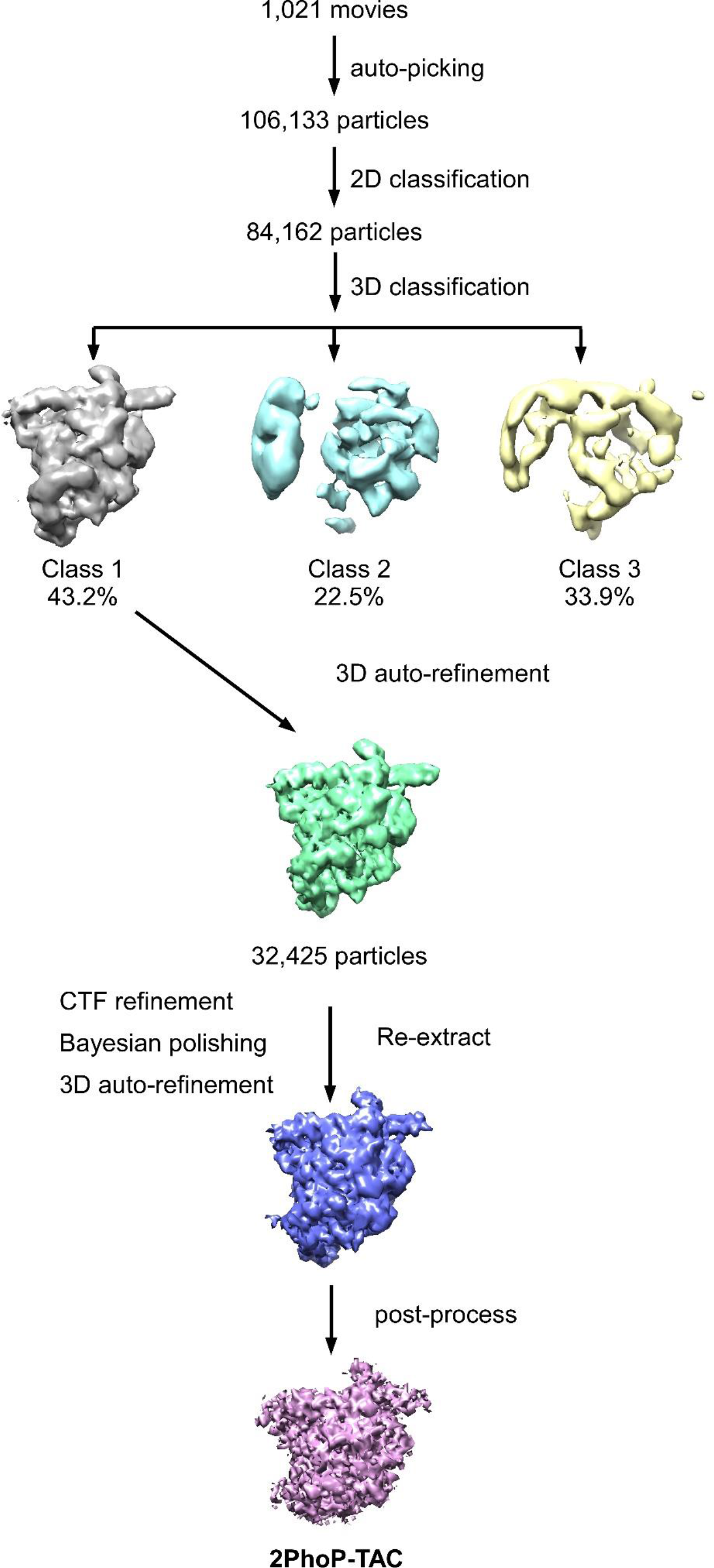
**Data processing pipeline for *Mtb* 2PhoP-TAC.**

**Supplementary Fig. 5.**
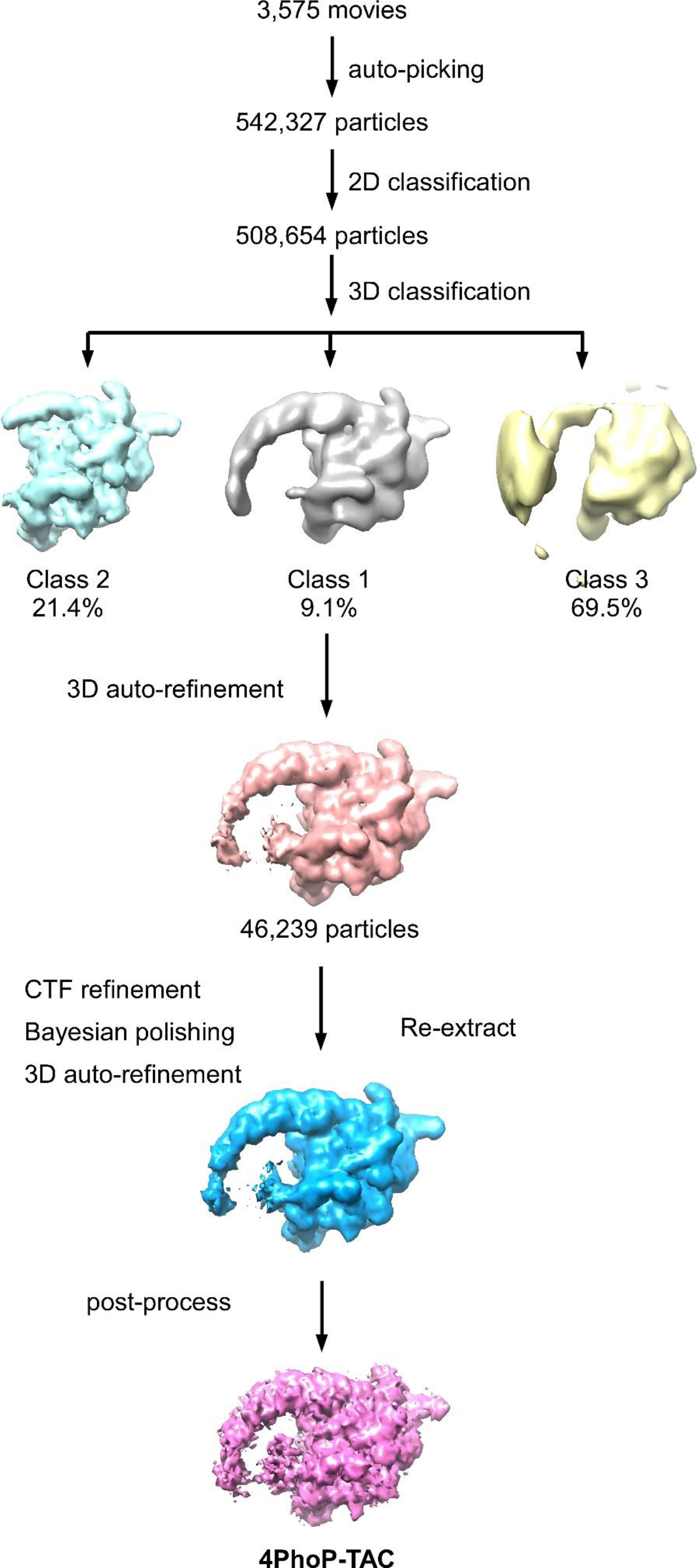
**Data processing pipeline for *Mtb* 4PhoP-TAC.**

**Supplementary Fig. 6.**
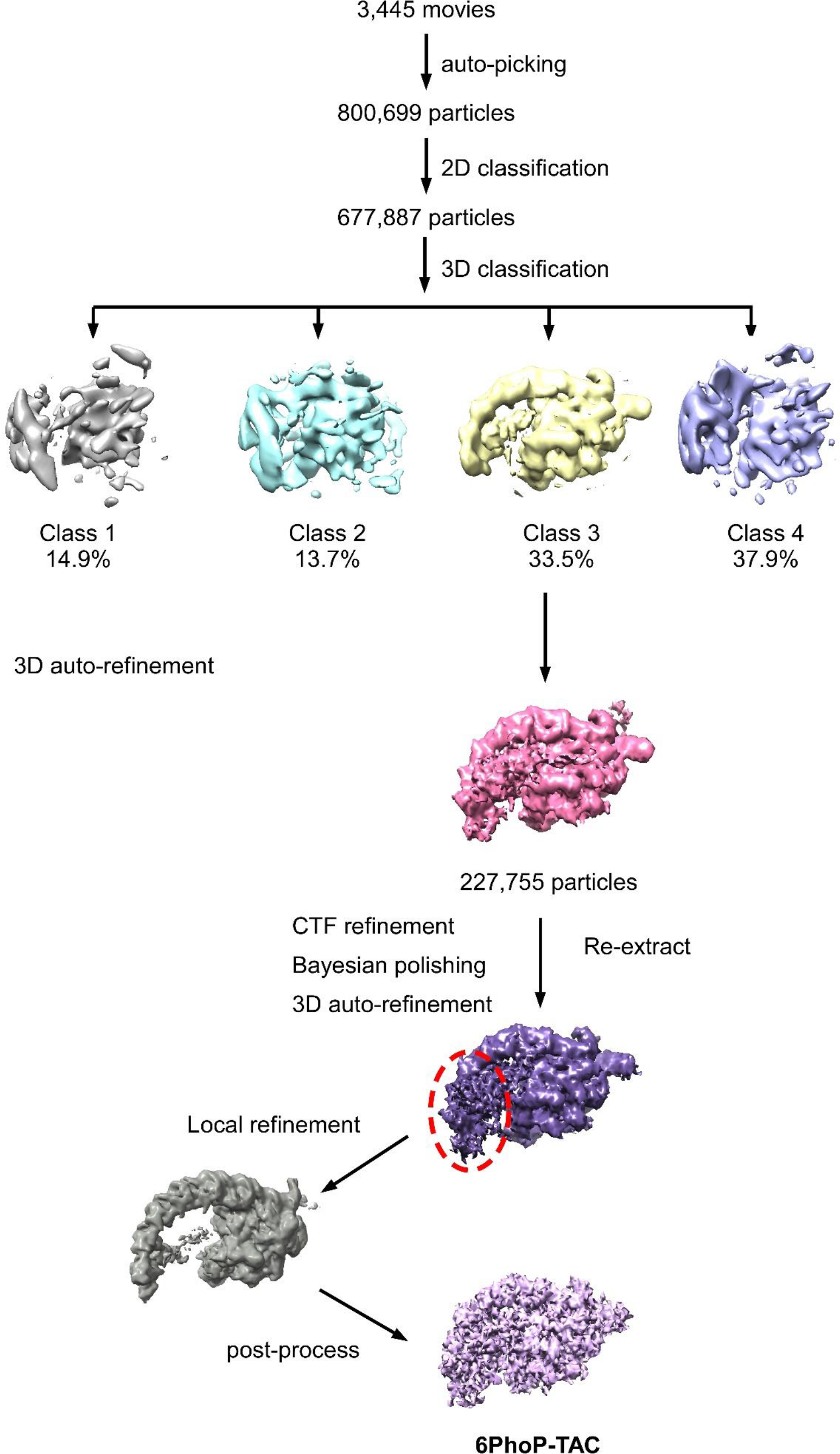
**Data processing pipeline for *Mtb* 6PhoP-TAC.**

**Supplementary Fig. 7.**
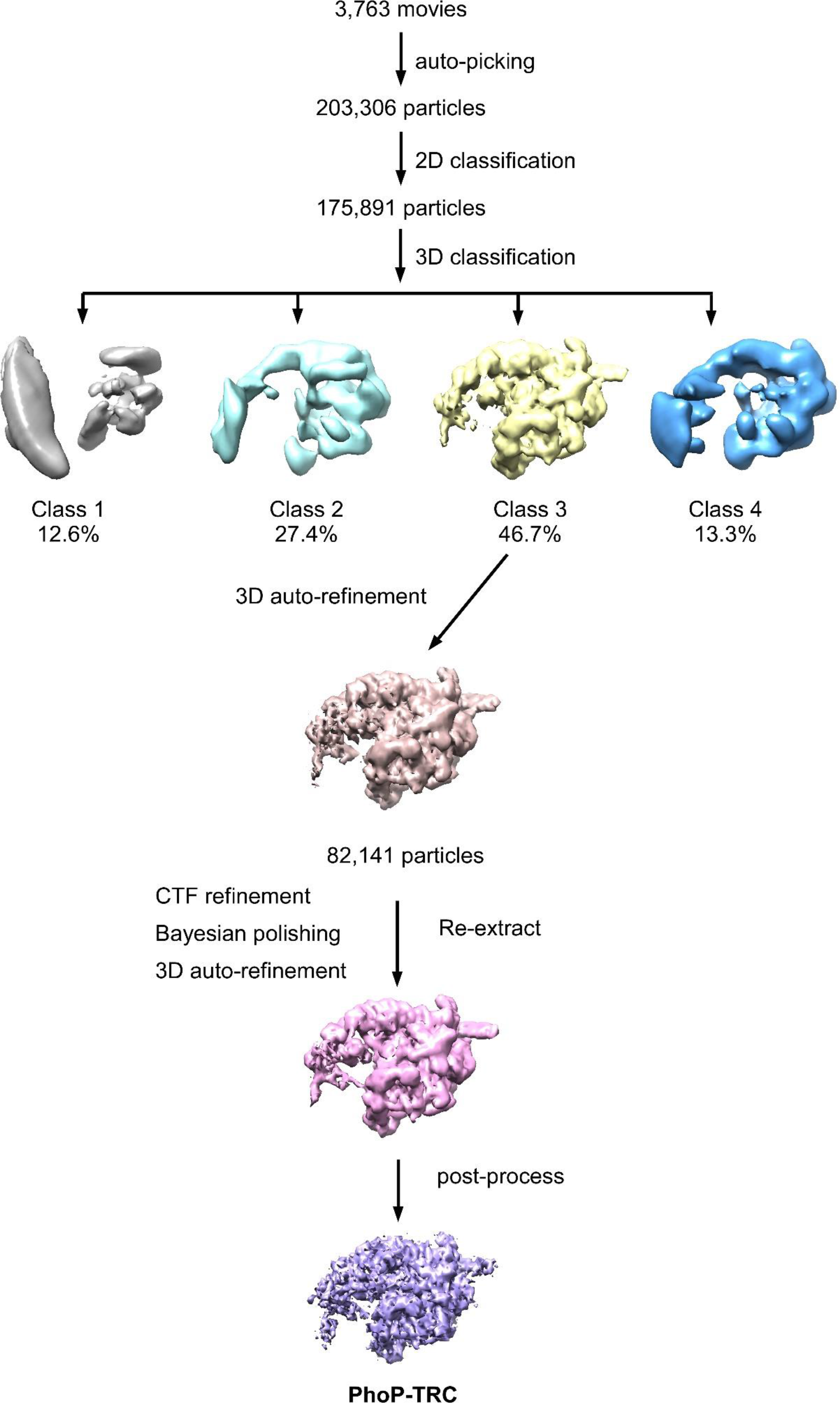
**Data processing pipeline for *Mtb* PhoP-TRC.**

**Supplementary Fig. 8.**
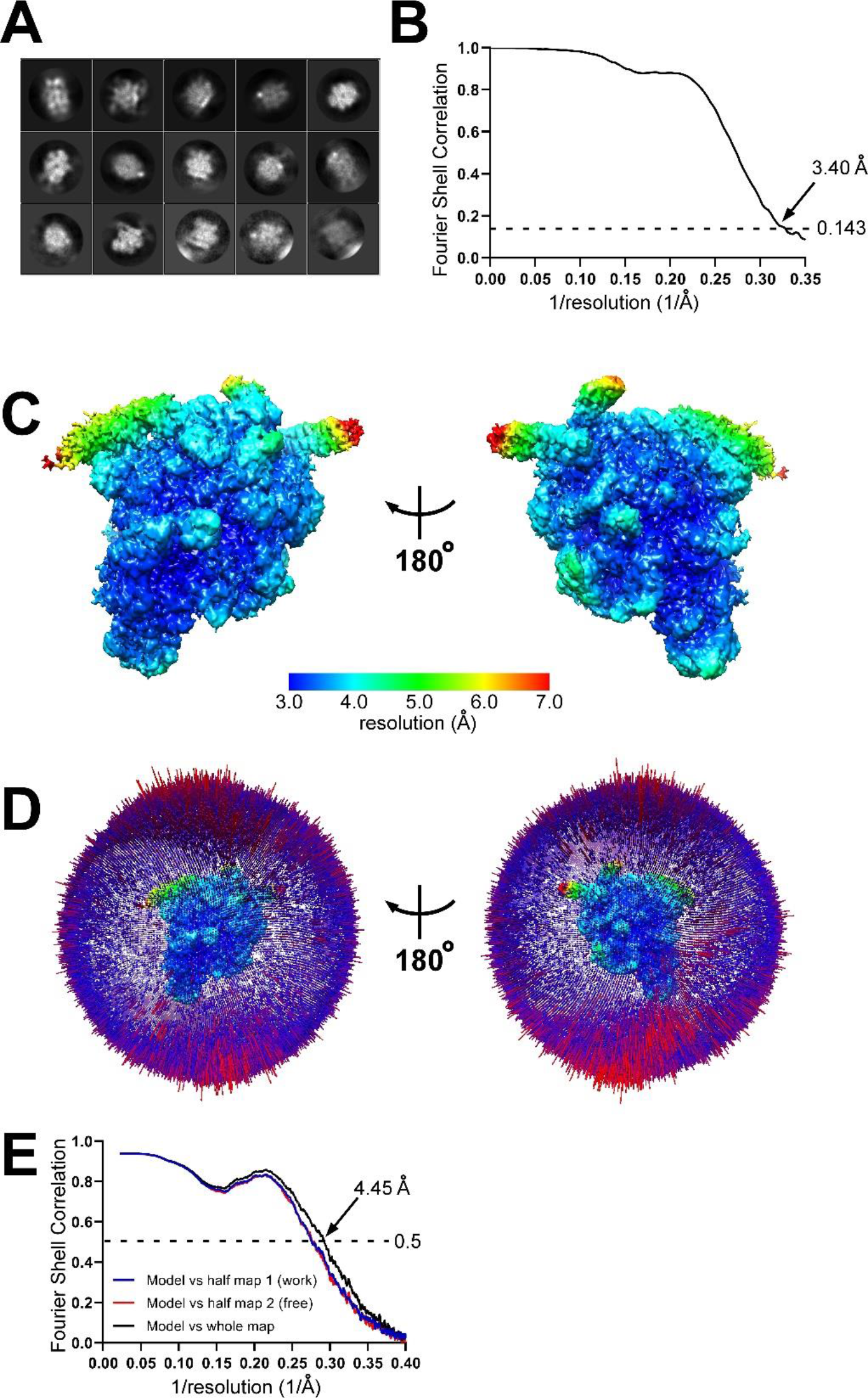
Cryo-EM data of *Mtb* RPo. a, Representative classes from 2D classification. **b,** Gold-standard FSC. The gold-standard FSC was calculated by comparing the two independently determined half-maps from RELION. The dashed line represents the 0.143 FSC cutoff ^1^, which indicates a nominal resolution of 3.40 Å. **c,** Cryo-EM density map colored by local resolution. Local resolution calculation was performed using blocres ^2^. View orientation as in Figure 1b. **d,** Angular distribution of particle projections. View orientations as in (c). **e,** FSC calculated between the model and the half map used for refinement (work), the other half map (free), and the full map.

**Supplementary Fig. 9.**
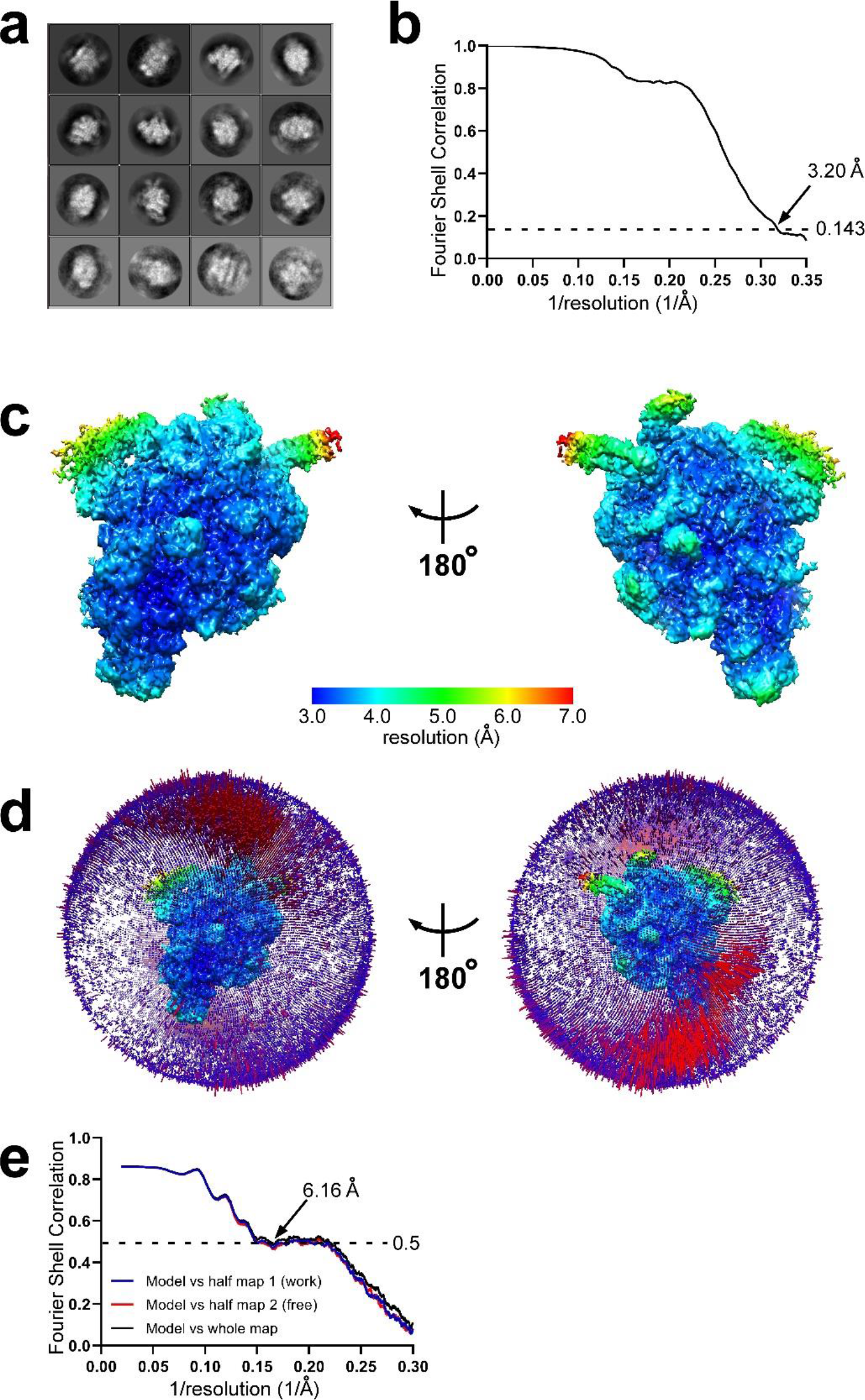
Cryo-EM data of *Mtb* 2PhoP-TAC. a, Representative classes from 2D classification. **b,** Gold-standard FSC. The gold-standard FSC was calculated by comparing the two independently determined half-maps from RELION. The dashed line represents the 0.143 FSC cutoff ^1^, which indicates a nominal resolution of 3.20 Å. **c,** Cryo-EM density map colored by local resolution. Local resolution calculation was performed using blocres ^2^. View orientation as in Figure 1b. **d,** Angular distribution of particle projections. View orientations as in (c). **e,** FSC calculated between the model and the half map used for refinement (work), the other half map (free), and the full map. **Supplementary** Fig.12 **Cryo-EM data of *Mtb* PhoP-TRC.**

**Supplementary Fig. 10.**
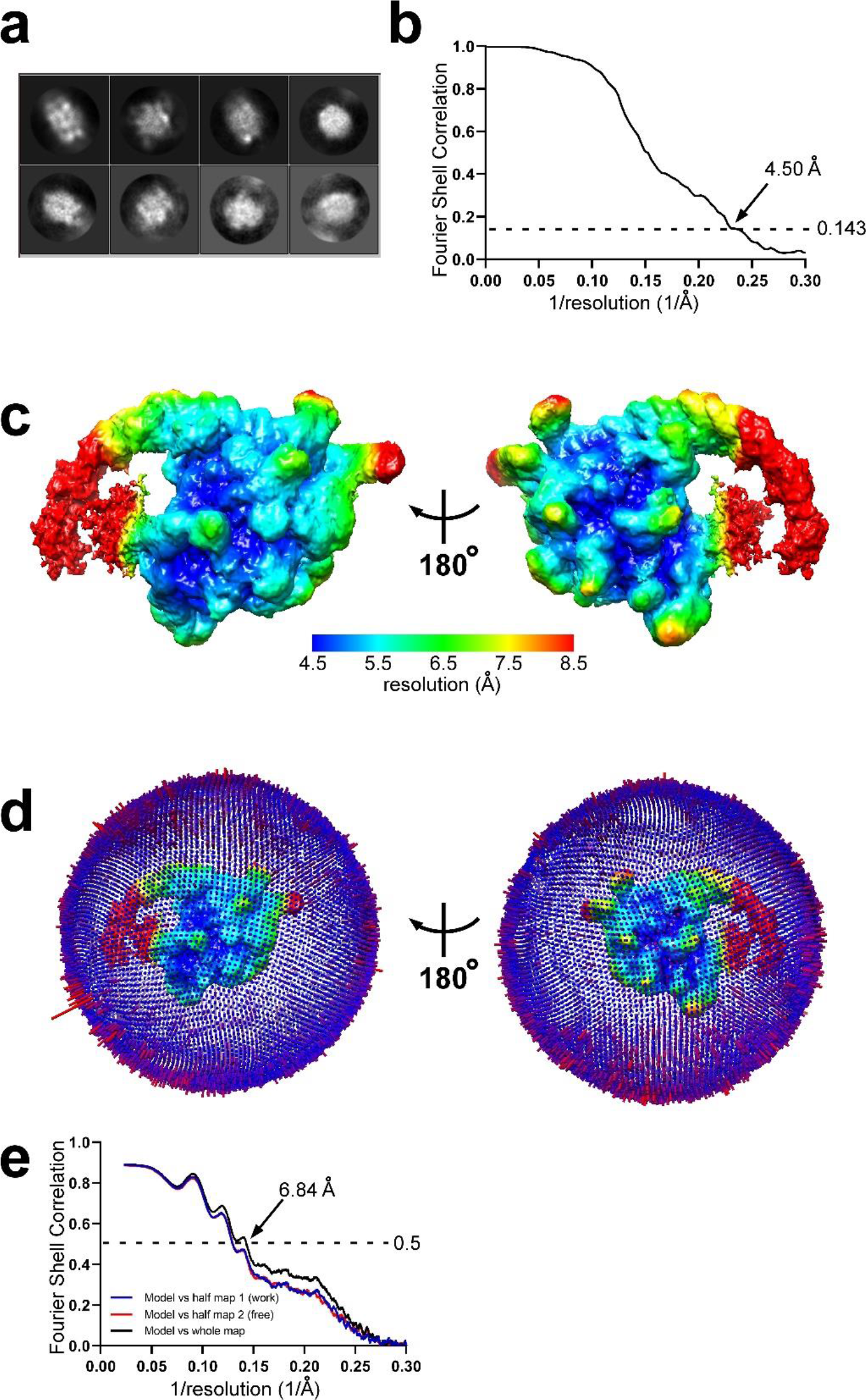
Cryo-EM data of *Mtb* 4PhoP-TAC. a, Representative classes from 2D classification. **b,** Gold-standard FSC. The gold-standard FSC was calculated by comparing the two independently determined half-maps from RELION. The dashed line represents the 0.143 FSC cutoff ^1^, which indicates a nominal resolution of 4.50 Å. **c,** Cryo-EM density map colored by local resolution. Local resolution calculation was performed using blocres ^2^. View orientation as in Figure 1b. **d,** Angular distribution of particle projections. View orientations as in (c). **e,** FSC calculated between the model and the half map used for refinement (work), the other half map (free), and the full map.

**Supplementary Fig. 11.**
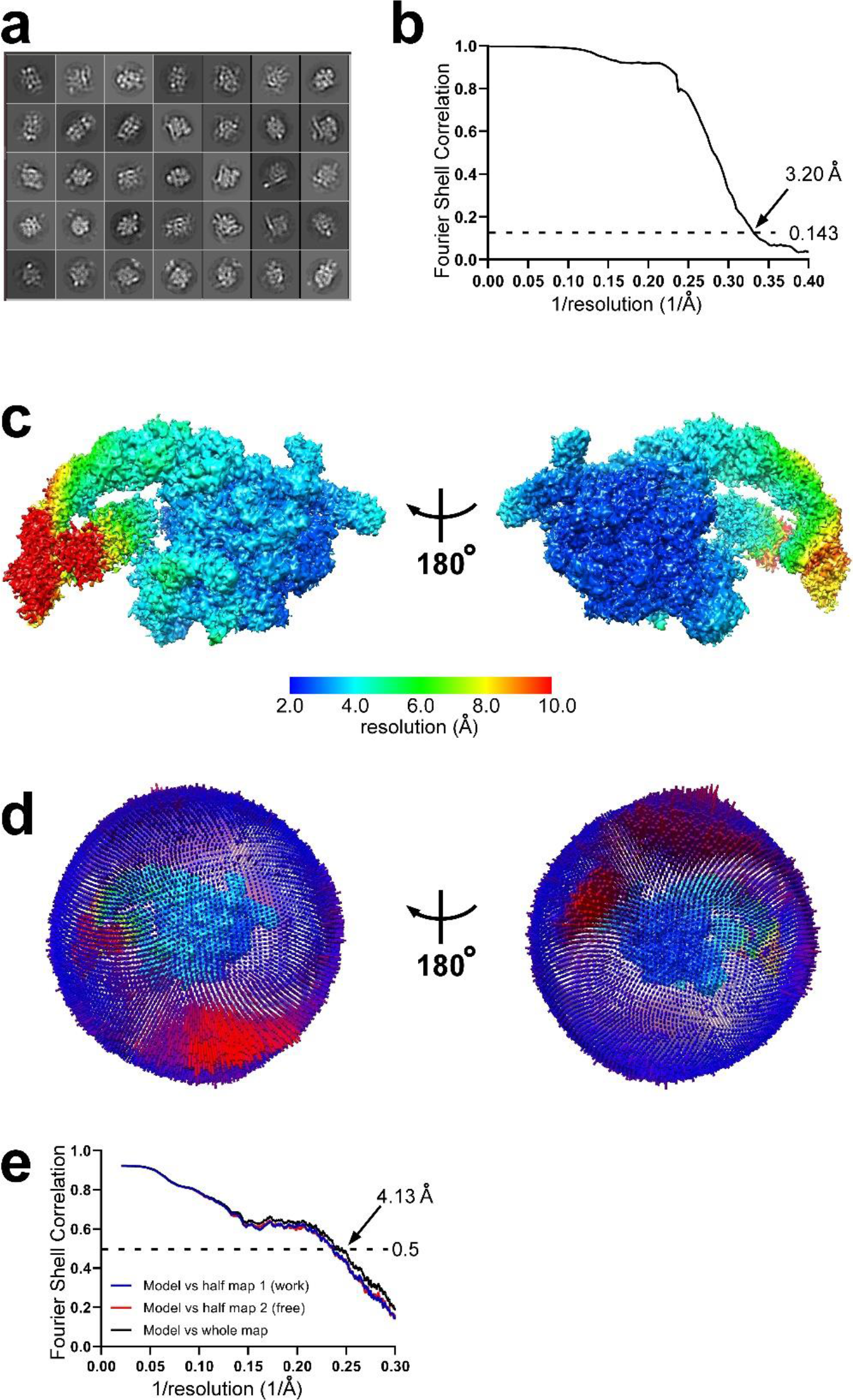
Cryo-EM data of *Mtb* 6PhoP-TAC. a, Representative classes from 2D classification. **b,** Gold-standard FSC. The gold-standard FSC was calculated by comparing the two independently determined half-maps from RELION. The dashed line represents the 0.143 FSC cutoff ^1^, which indicates a nominal resolution of 3.20 Å. **c,** Cryo-EM density map colored by local resolution. Local resolution calculation was performed using blocres ^2^. View orientation as in Figure 1b. **d,** Angular distribution of particle projections. View orientations as in (c). **e,** FSC calculated between the model and the half map used for refinement (work), the other half map (free), and the full map.

**Supplementary Fig. 12.**
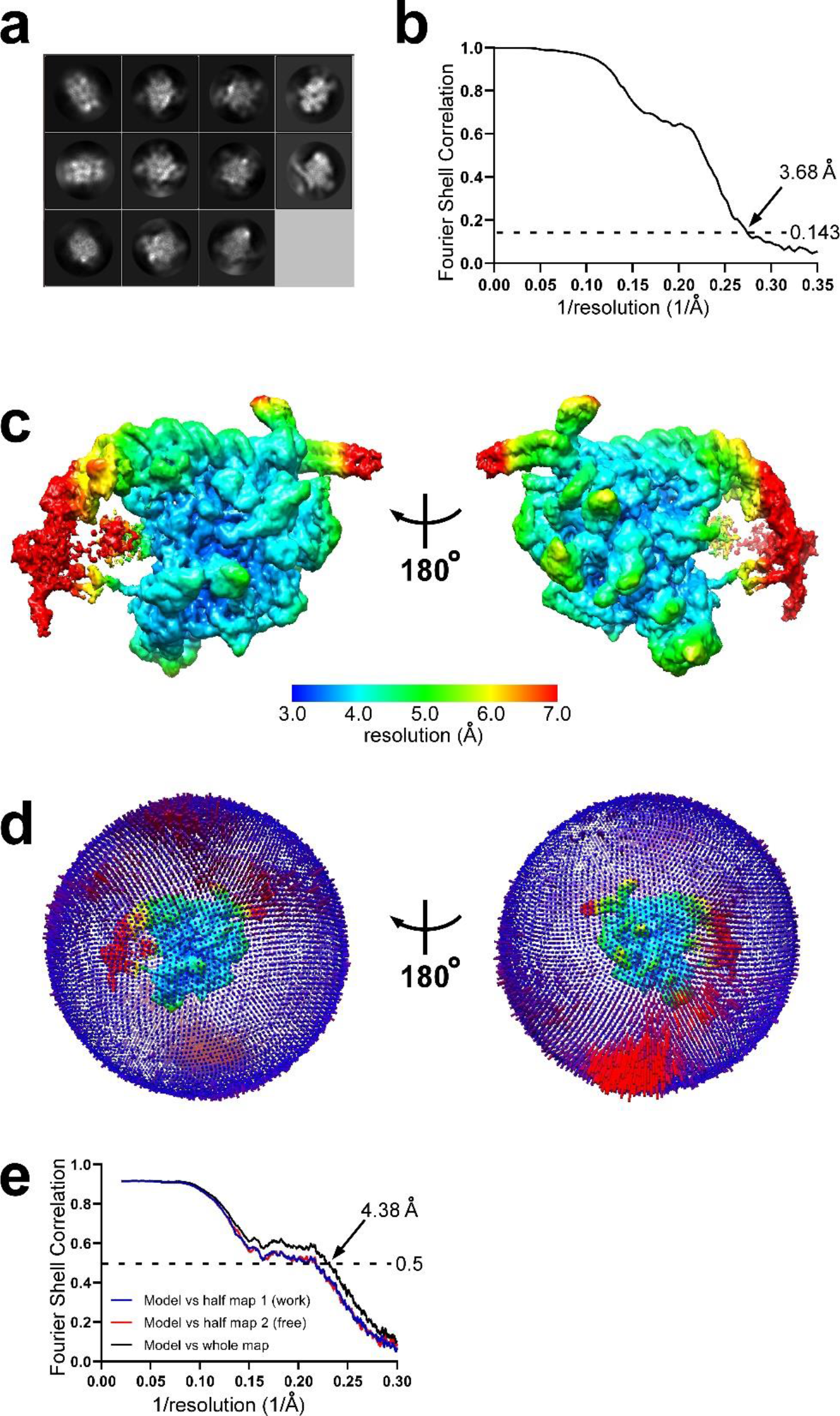
Cryo-EM data of *Mtb* PhoP-TRC. a, Representative classes from 2D classification. **b,** Gold-standard FSC. The gold-standard FSC was calculated by comparing the two independently determined half-maps from RELION. The dashed line represents the 0.143 FSC cutoff ^1^, which indicates a nominal resolution of 3.68 Å. **c,** Cryo-EM density map colored by local resolution. Local resolution calculation was performed using blocres ^2^. View orientation as in Figure 1b. **d,** Angular distribution of particle projections. View orientations as in (c). **e,** FSC calculated between the model and the half map used for refinement (work), the other half map (free), and the full map.

**Supplementary Fig. 13.**
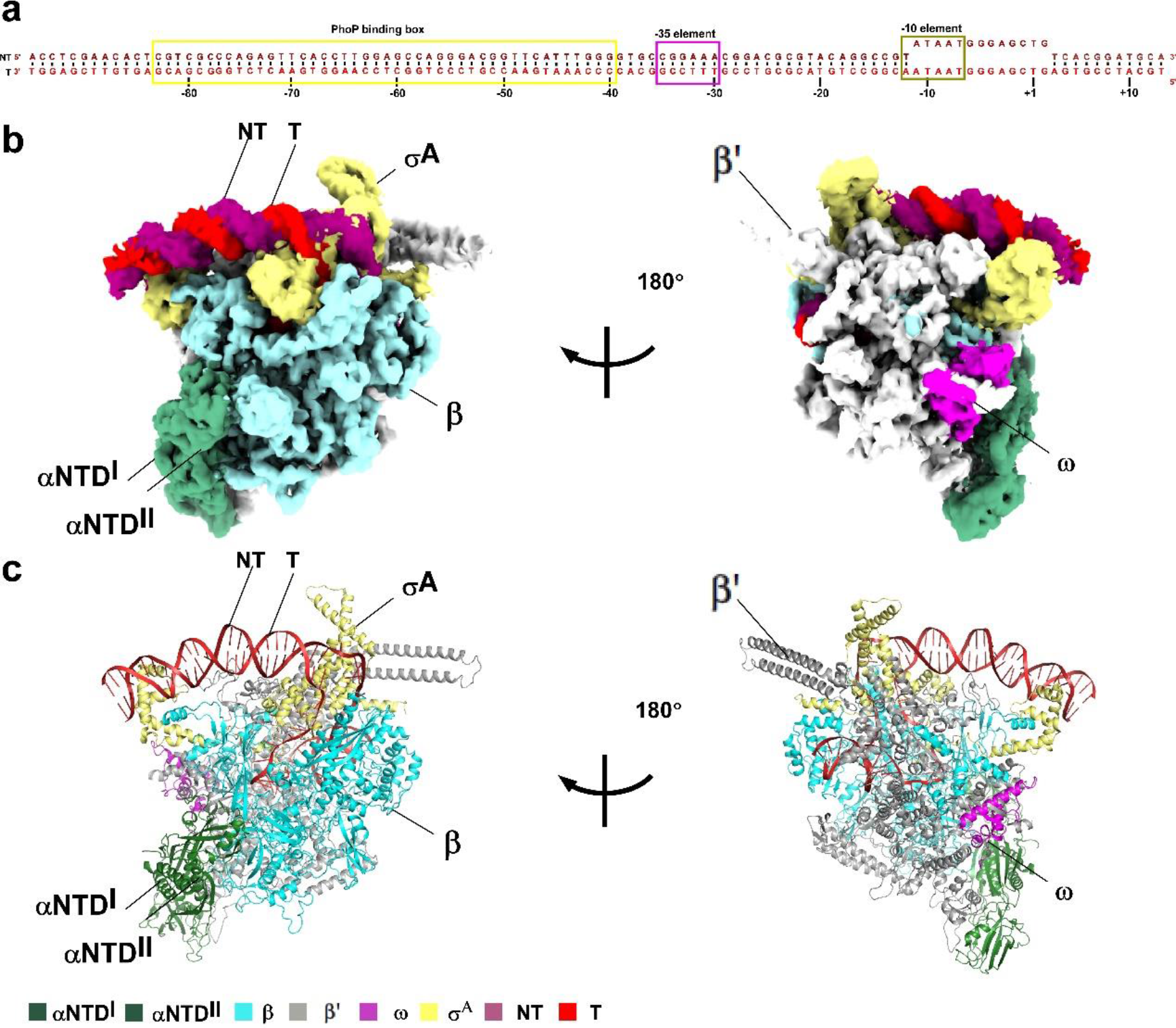
The overall structure of *Mtb* RPo. a, DNA scaffold used in structure determination of *Mtb* RPo. PhoP binding box is framed in yellow color. **b- c,** Two views of the cryo-EM density map **(b)** and structure model **(c)** of *Mtb* RPo. The EM density map is colored as indicated in the color key.

**Supplementary Fig. 14.**
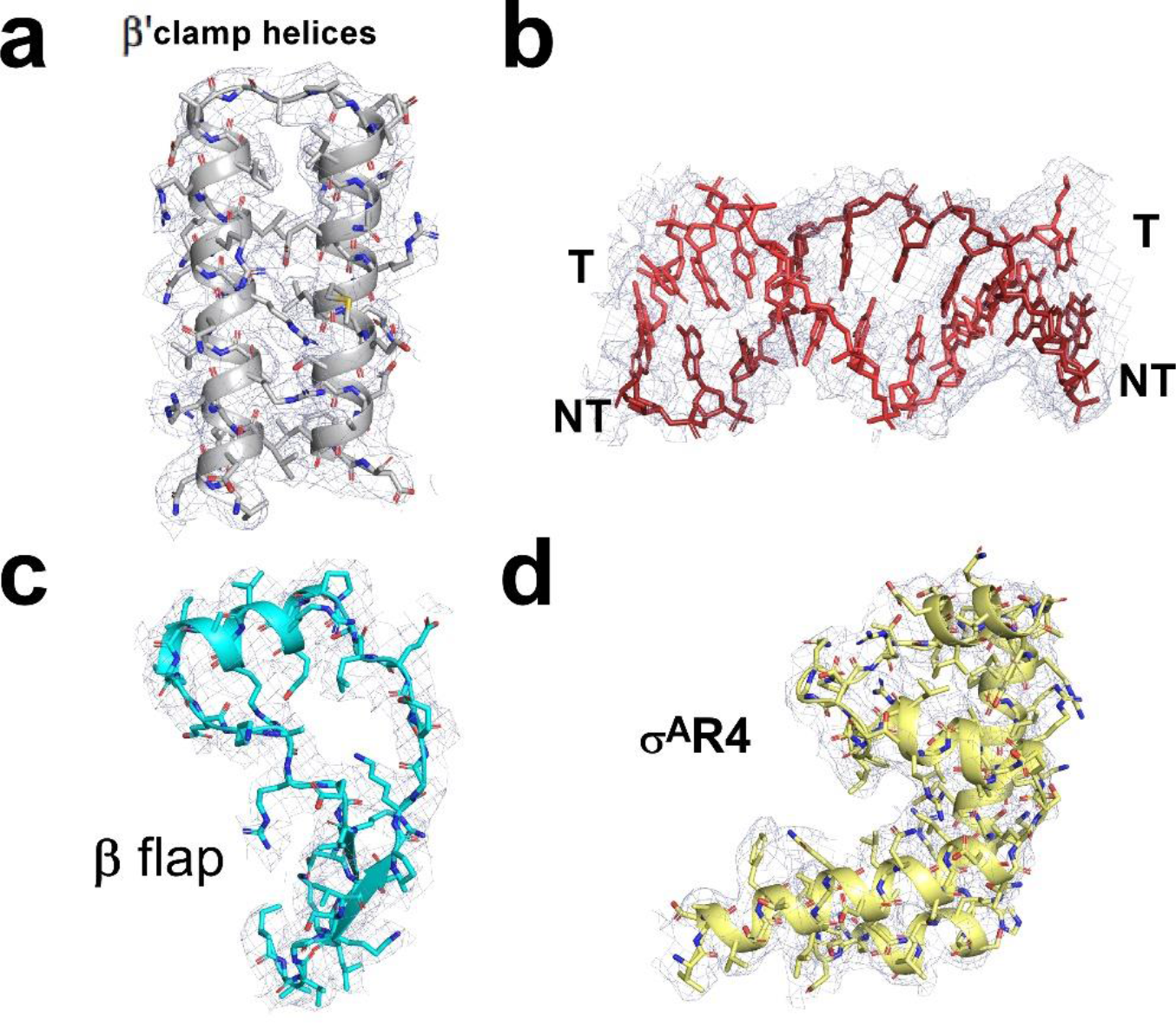
Representative cryo-EM densities of superimposed models in *Mtb* RPo. **a,** Cryo-EM density map (grey mesh) and the superimposed model of β’ clamp helices. **b,** Cryo-EM density map (grey mesh) and the superimposed model of PhoP binding site. **c,** Cryo-EM density map (grey mesh) and the superimposed model of RNAP β flap. **d,** Cryo-EM density map (grey mesh) and the superimposed model of σ^A^R4. Other colors are shown as in **Supplementary** Fig.13.

**Supplementary Fig. 15.**
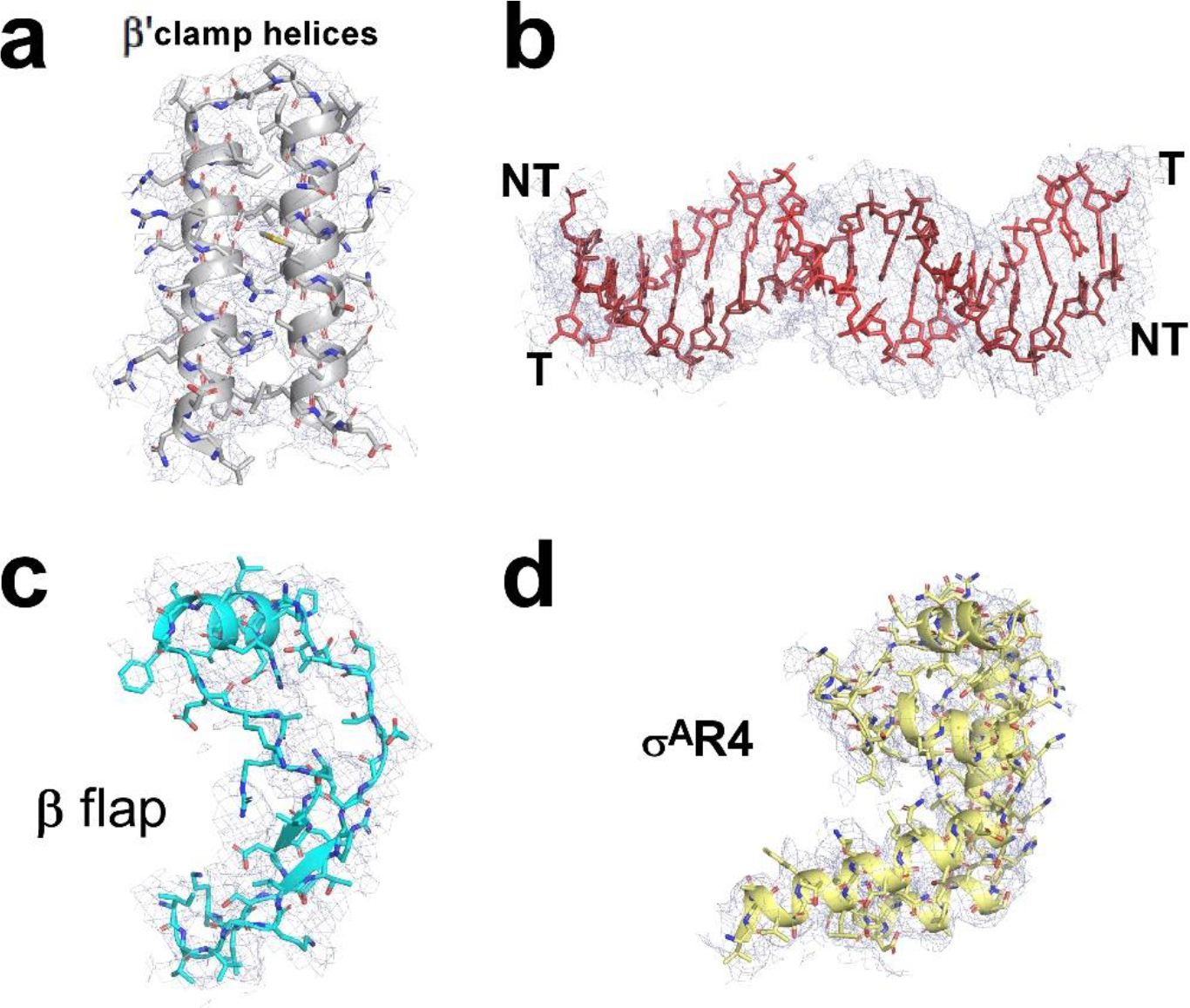
Representative cryo-EM densities of superimposed models in *Mtb* 2PhoP-TAC. **a,** Cryo-EM density map (grey mesh) and the superimposed model of β’ clamp helices. **b,** Cryo-EM density map (grey mesh) and the superimposed model of PhoP binding site. **c,** Cryo-EM density map (grey mesh) and the superimposed model of RNAP β flap. **d,** Cryo-EM density map (grey mesh) and the superimposed model of σ^A^R4. Other colors are shown as in **Fig. 1a**.

**Supplementary Fig. 16.**
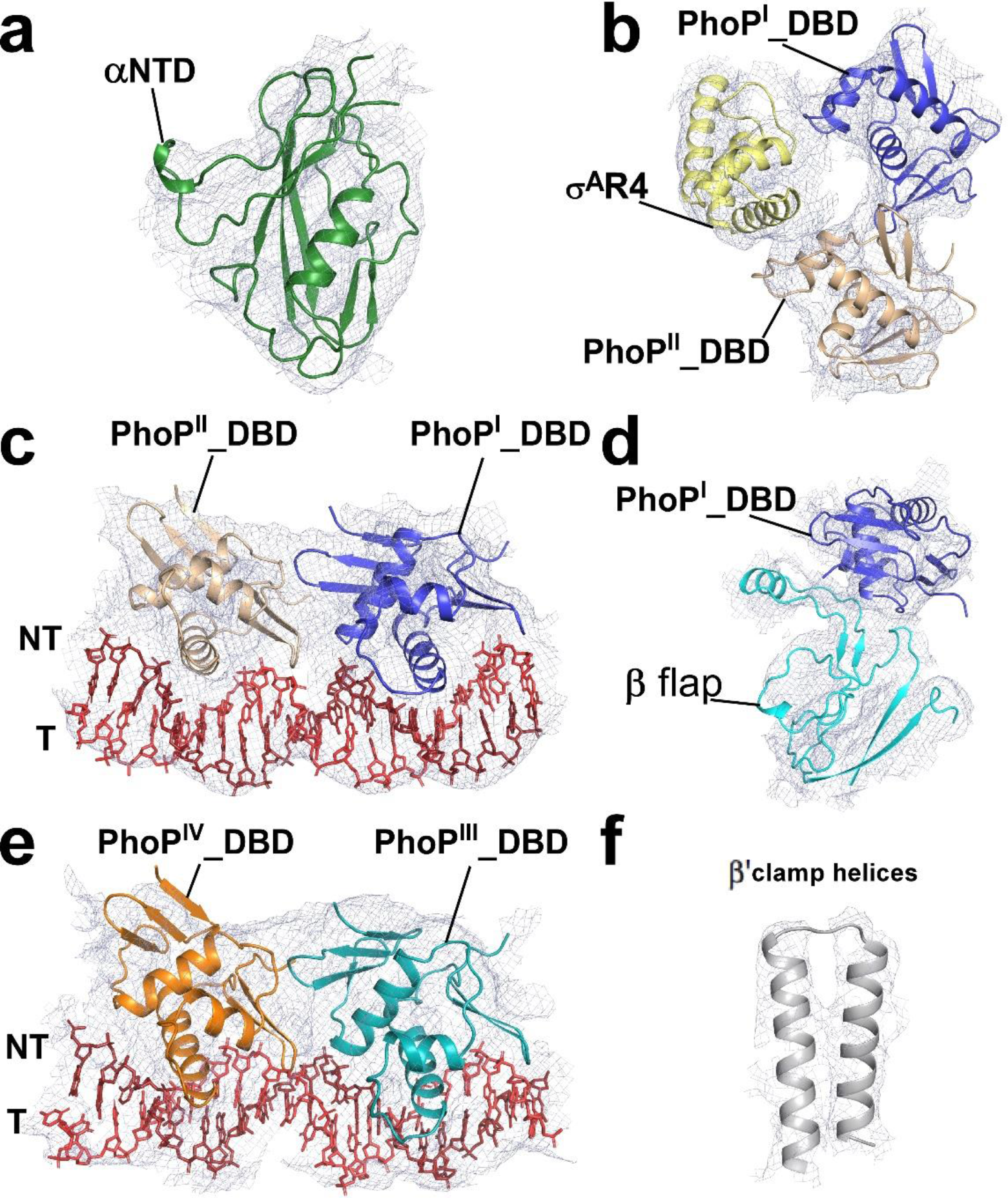
Representative cryo-EM densities of superimposed models in *Mtb* 4PhoP-TAC. **a,** Cryo-EM density map (grey mesh) and the superimposed model of RNAP αNTD. **b,** Cryo-EM density map (grey mesh) and the superimposed model of PhoP^I^_DBD PhoP^II^_DBD and σ^A^R4. **c,** Cryo-EM density map (grey mesh) and the superimposed model of PhoP^I^_DBD PhoP^II^_DBD and their PhoP binding sites. **d,** Cryo-EM density map (grey mesh) and the superimposed model of PhoP^I^_DBD and β flap. **e,** Cryo-EM density map (grey mesh) and the superimposed model of PhoP^IV^_DBD PhoP^III^_DBD and their PhoP binding site. **f,** Cryo-EM density map (grey mesh) and the superimposed model of β’ clamp helices. Other colors are shown as in **Fig. 1b**.

**Supplementary Fig. 17.**
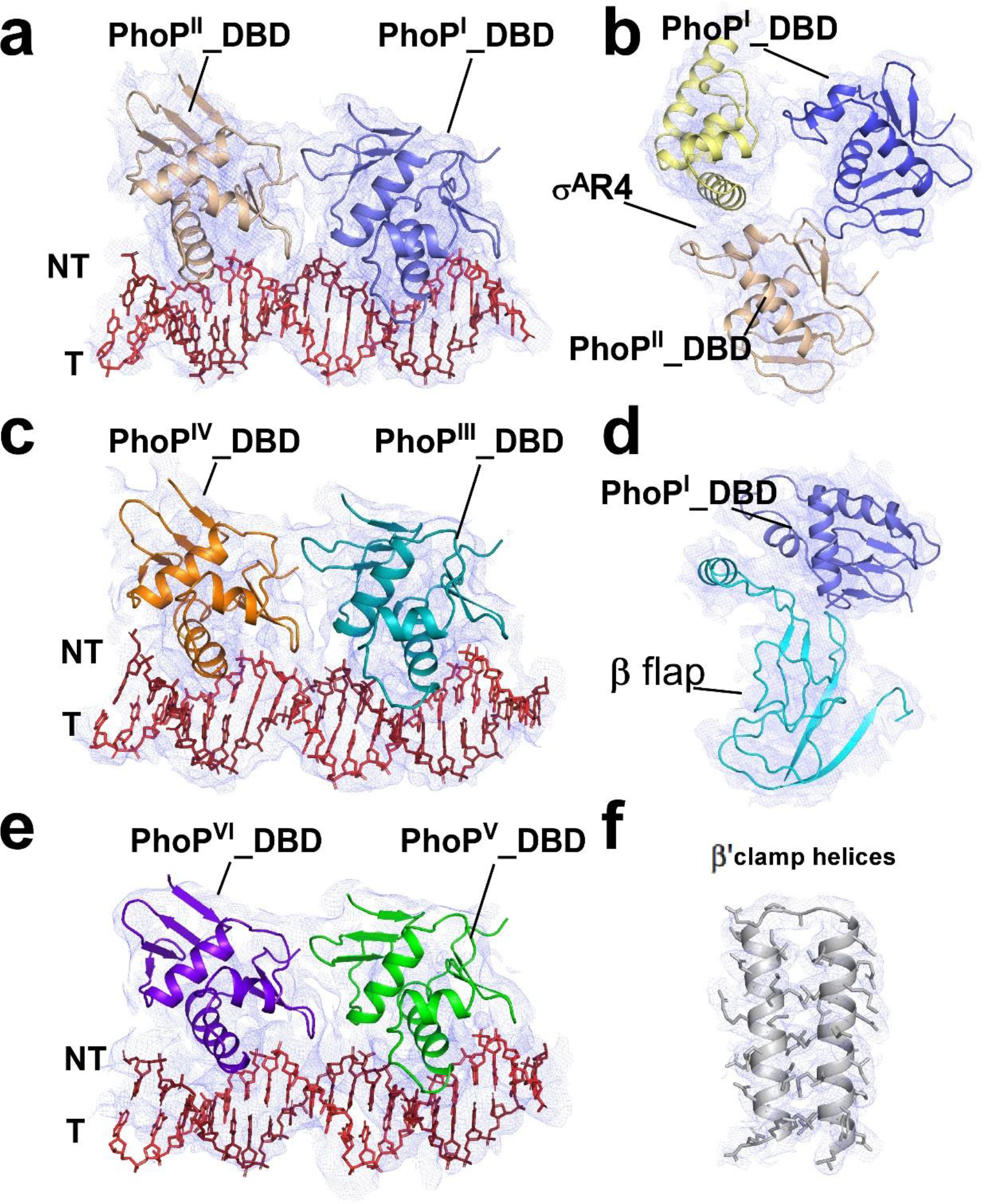
Representative cryo-EM densities of superimposed models in *Mtb* 6PhoP-TAC. **a,** Cryo-EM density map (grey mesh) and the superimposed model of PhoP^I^_DBD PhoP^II^_DBD and their PhoP binding sites. **b,** Cryo-EM density map (grey mesh) and the superimposed model of PhoP^I^_DBD PhoP^II^_DBD and σ^A^R4. **c,** Cryo-EM density map (grey mesh) and the superimposed model of PhoP^III^_DBD PhoP^IV^_DBD and their PhoP binding sites. **d,** Cryo-EM density map (grey mesh) and the superimposed model of PhoP^I^_DBD and β flap. **e,** Cryo-EM density map (grey mesh) and the superimposed model of PhoP^V^_DBD PhoP^VI^_DBD and their PhoP binding site. **f,** Cryo-EM density map (grey mesh) and the superimposed model of β’ clamp helices. Other colors are shown as in **Fig. 1c**.

**Supplementary Fig. 18.**
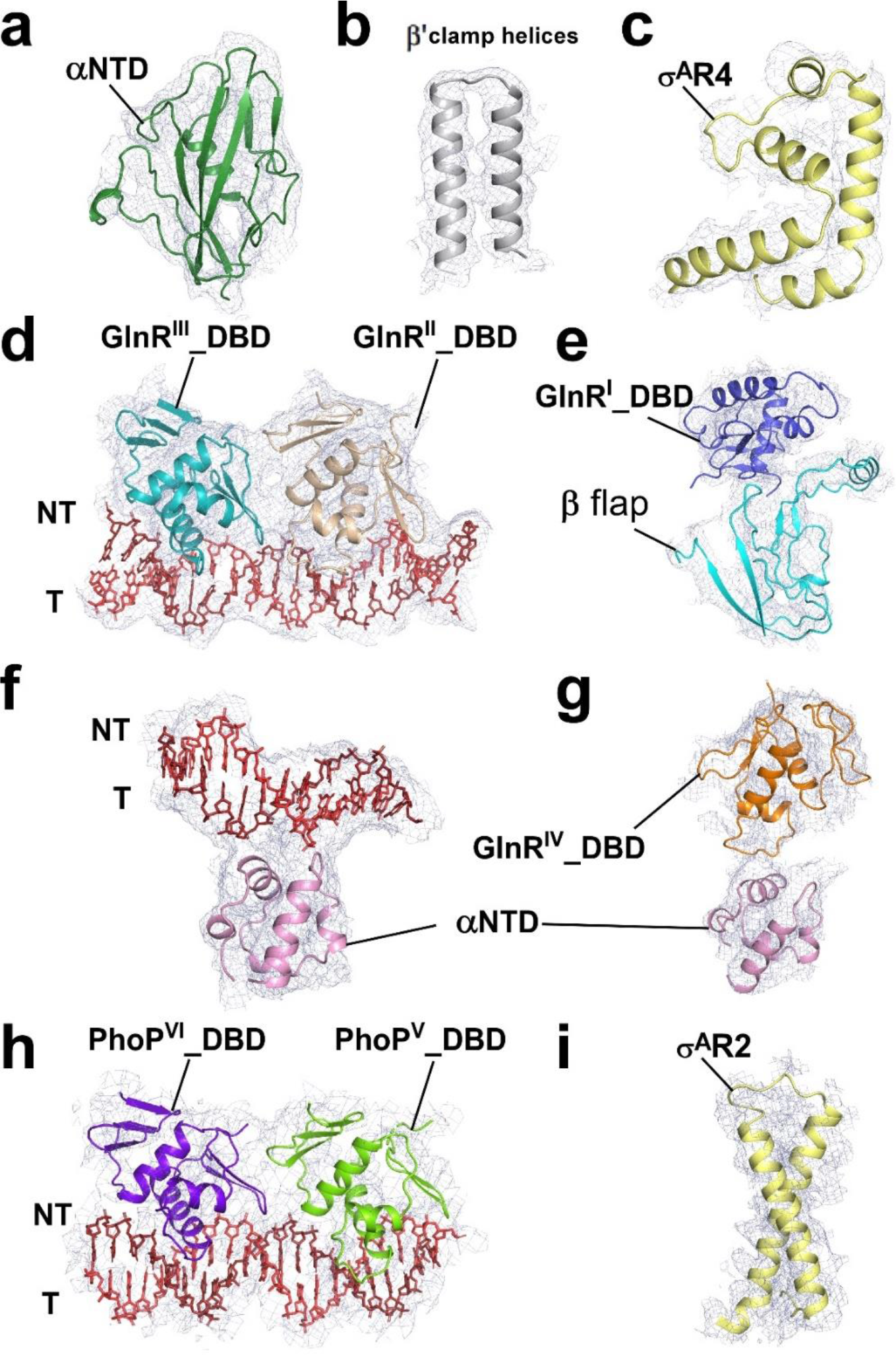
Representative cryo-EM densities of superimposed models in *Mtb* PhoP-TRC. **a,** Cryo-EM density map (grey mesh) and the superimposed model of RNAP αNTD. **b,** Cryo-EM density map (grey mesh) and the superimposed model of β’ clamp helices. **c,** Cryo-EM density map (grey mesh) and the superimposed model of σ^A^R4. **d,** Cryo-EM density map (grey mesh) and the superimposed model of GlnR^II^_DBD, GlnR^III^_DBD and their GlnR binding sites. **e,** Cryo-EM density map (grey mesh) and the superimposed model of GlnR^I^_DBD and β flap. **f-g,** Cryo-EM density map and the superimposed model of αCTD and the promoter DNA (**f**) and of αCTD and GlnR^IV^_DBD (**g**). **h,** Cryo-EM density map (grey mesh) and the superimposed model of PhoP^V^_DBD, PhoP^VI^_DBD and their PhoP binding sites. **i,** Cryo-EM density map (grey mesh) and the superimposed model of σ^A^R2. Other colors are shown as in **Fig. 4c**.

**Supplementary Fig. 19.**
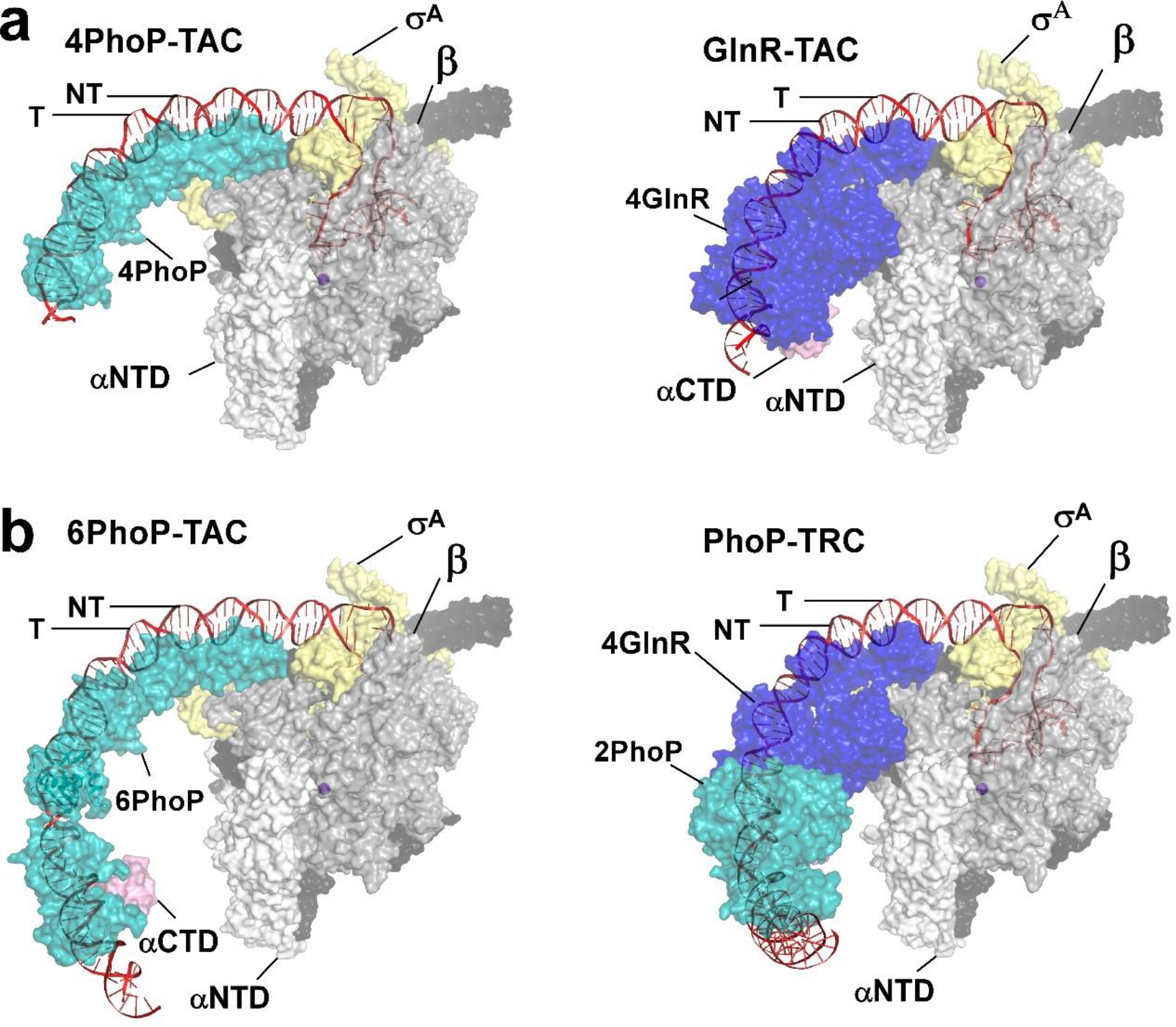
Structural comparisons of *Mtb* 4PhoP-TAC, *Mtb* GlnR-TAC, *Mtb* 6PhoP-TAC, and *Mtb* PhoP-TRC **a,** *Mtb* 4PhoP-TAC (left panel) and *Mtb* GlnR-TAC (right panel, PDB ID: 8HIH) ^3^ are shown in surface. **b,** *Mtb* 6PhoP-TAC (left panel) (left panel) and *Mtb* PhoP-TRC (right panel) are shown in surface. Cyan, PhoP; blue, GlnR; yellow, σ^A^; white, pink, wheat, gray, and dark gray, RNAP αNTD, αCTD, β and β’. Other colors are shown as in **Fig. 1c** and **Fig. 4b**.

**Supplementary Table S1.**
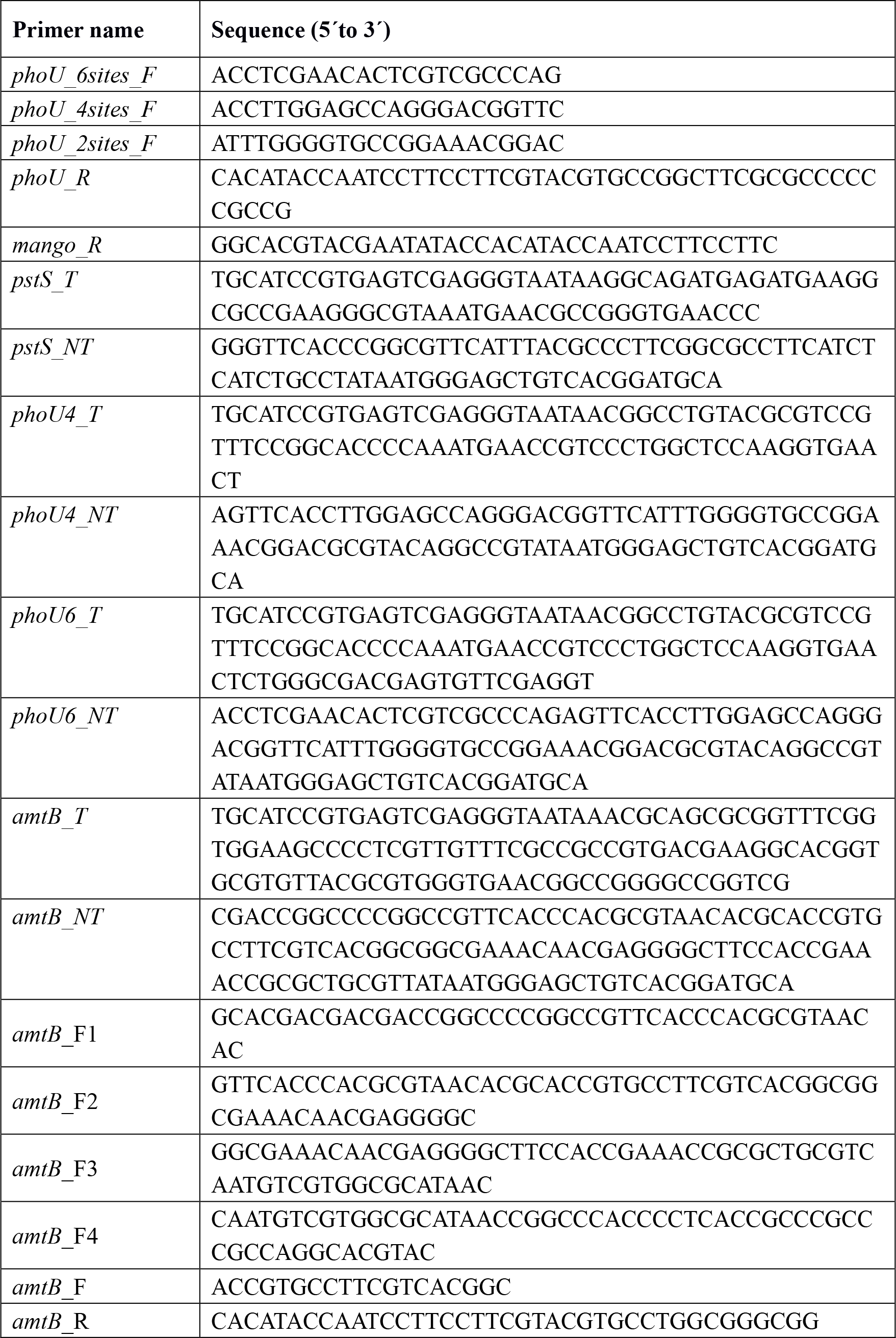
Primers used in this study.

